# Start codon context controls translation initiation in the fungal kingdom

**DOI:** 10.1101/654046

**Authors:** Edward Wallace, Corinne Maufrais, Jade Sales-Lee, Laura Tuck, Luciana de Oliveira, Frank Feuerbach, Frédérique Moyrand, Prashanthi Natarajan, Hiten D. Madhani, Guilhem Janbon

## Abstract

Eukaryotic protein synthesis initiates at a start codon defined by an AUG and its surrounding Kozak sequence context, but studies of S. *cerevisiae* suggest this context is of little importance in fungi. We tested this concept in two pathogenic *Cryptococcus* species by genome-wide mapping of translation and of mRNA 5’ and 3’ ends. We observed that upstream open reading frames (uORFs) are a major contributor to translation repression, that uORF use depends on the Kozak sequence context of its start codon, and that uORFs with strong contexts promote nonsense-mediated mRNA decay. Numerous *Cryptococcus* mRNAs encode predicted dual-localized proteins, including many aminoacyl-tRNA synthetases, in which a leaky AUG start codon is followed by a strong Kozak context in-frame AUG, separated by mitochondrial-targeting sequence. Further analysis shows that such dual-localization is also predicted to be common in *Neurospora crassa*. Kozak-controlled regulation is correlated with insertions in translational initiation factors in fidelity-determining regions that contact the initiator tRNA. Thus, start codon context is a signal that programs the expression and structures of proteins in fungi.

## Introduction

Fungi are important in the fields of ecology, medicine, and biotechnology. With 4 million predicted fungal species, this kingdom is the most diverse of the domain Eukarya. Recent initiatives such as the 1000 Fungal Genomes Project at the Joint Genome Institute, or the Global Catalogue of Microorganisms, which aims to produce 2500 complete fungal genomes in the next 5 years, will result in a deluge of genome sequence data (1, 2). Comparative analysis of coding sequences enables the generation of hypotheses on genome biology and evolution (3–6). However, these analyses intrinsically depend on the quality of the coding gene identification and annotation, which have limitations. First, they depend on automatic sequence comparisons, which limit the identification of clade-specific genes. Second, fungal genes generally contain introns whose positions are difficult to predict based on the genome sequence alone (7). An uncertain intron annotation results in a poor annotation of the coding region extremities, which are generally less evolutionary conserved (8). Third, annotation pipelines only predict plausible open reading frames (ORFs), initially for yeast a contiguous stretch of at least 100 codons starting with an AUG codon and ending with a stop codon (9). These approaches do not reveal which ORFs are translated to protein, and are biased against short ORFs (10).

The common assumption is that the first AUG of a fungal ORF is used as the translation start codon, yet non-AUG start codons have been observed in every studied eukaryote, including the fungi S. *cerevisiae, C. albicans, S. pombe*, and *N. crassa* (11–16). In metazoans the rules for selection of AUG start codons were discovered by Kozak: AUGs are efficiently selected by mammalian cells if they are far (>20nt) from the transcription start site, and have a strong sequence context gccRccAUGG, where the R indicates a purine at the −3 position (17, 18). The influence of the motif on the efficiency of translation is organism-dependent. Although in metazoan cells the Kozak context has a major effect on translation initiation (19), in S. *cerevisiae* translation usually starts at the first AUG in the mRNA sequence, the strong aaaAUG sequence context only weakly affects endogenous protein output (20), and AUG sequence context somewhat modulates protein output from reporter mRNAs (21, 22). In fact, translation start codon context has a larger effect on non-AUG start codon usage (23). These *“Saccharomyces* rules” have been considered as the paradigm for fungi, but relevant data are lacking in other species of fungi.

Weak or inefficient start codons near the 5’end of mRNA can give rise to translational regulation, explained by the scanning model of eukaryotic translation initiation. Translation starts by the pre-initiation complex binding mRNA at the 5’ cap and then scanning the transcript leader (TL) sequence until it identifies a start codon, at which translation initiates (19). Here we call the 5’ regulatory region of mRNA the TL rather than the 5’ UTR, because short “upstream” ORFs in this region can be translated (24). The pre-initiation complex sediments at 43S, and comprises the small ribosomal subunit, methionyl initiator tRNA, and numerous eukaryotic translation initiation factors (eIFs). Biochemical, genetic, and structural data indicate that eIF1 and eIF1A associate with the 43S pre-initiation complex (25, 26). Recognition of the start codon involves direct interactions of eIF1 and eIF1A with the start codon context and initiator tRNA within a larger eIF2/3/5-containing 48S pre-initiation complex. Start codon selection occurs when eIF1 is replaced by eIF5’s N-terminus (27), then eIF2 is released, the large ribosomal subunit joins catalyzed by eIF5B, and translation begins (26). This work has been largely driven by studies in S. *cerevisiae* and metazoans. Although the core protein and RNA machinery of eukaryotic translation initiation is highly conserved, it is not understood how fungi quantitatively vary in the sequence, structure, and function of their translation initiation machinery.

*Cryptococcus* are basidiomycete yeasts with a high density of introns in their coding genes (28). These introns influence gene expression and genome stability (29–31). The current genome annotation of pathogenic *C. neoformans* and *deneoformans* reference strains are based on both automatic and manual curations of gene structures using RNA-Seq data (32, 33). Although the high degree of interspecies conservation of intron numbers and positions within coding sequences suggest that these annotations are reliable (33), the regulatory regions (transcript leader and 3’ UTRs) at transcript extremities are less well identified. In fact, most fungal genomes lack complete transcript annotations, and thus we do not know how regulatory structure varies across fungi.

In this paper, we experimentally determine the beginning and the end of both coding regions and of transcripts in two *Cryptococcus* species, providing an important genomic resource for the field. Furthermore, our joint analysis of TL sequences and translation identifies a Kozak sequence context that regulates start codon selection, affecting upstream ORF regulation and also alternative protein targeting to mitochondria. Comparison with other fungal genomes revealed that these types of regulation are common in this kingdom: the first AUG of an mRNA or an ORF is not always the major start codon in fungi. These studies demonstrate that, in contrast to the situation in S. *cerevisiae*, start codon sequence context is an important gene regulatory signal that programs the levels and structures of proteins in the fungal kingdom.

## RESULTS

### Delineation of transcript ends in *C. neoformans* and *C. deneoformans*

To annotate the extremities of the coding genes in *C. neoformans* and *C. deneoformans, we* mapped the 5’ ends (Transcription Start Sites; TSS) with TSS-Seq (34), the 3’ ends (Polyadenylation sites; PAS) with QuantSeq 3’mRNA-Seq, and sequenced the same samples with stranded mRNA-Seq. These experiments were done in biological triplicate from cells growing at two temperatures (30°C and 37°C) and two stages of growth (exponential and early stationary phases) with external normalization with spike-in controls.

We identified 4.7 × 10^6^ unique TSSs and 6.3 × 10^4^ unique PASs in *C. neoformans*. Clustering of these positions revealed between 27,339 and 42,720 TSS clusters and between 9,217 and 16,697 PAS clusters depending on the growth conditions (Table S1). We used the clusters associated with the coding genes to produce an initial annotation, using the most distal TSS and PAS clusters for each gene. The predicted positions which changed the extremities of the genes by more than 100 bp were manually curated (n=1131 and n=286 for the TSS and PAS, respectively). We then selected the most prominent clusters that represented at least 10% of the normalized reads count per coding gene in at least one condition. Finally, the most distal of these TL-TSS and 3’UTR-PA clusters were labeled as the 5’and 3’ ends of the coding genes for our final annotation (Table S1). For the genes for which no TL-TSS cluster or no 3’UTR-PAS cluster could be identified, we maintained the previous annotation. We used the same strategies for *C. deneoformans* and obtained similar results (Table S1).

As expected, most of the TSS clusters (62%) were associated with the TL whereas most of the PAS clusters (82%) were associated with the 3’UTR of the coding genes (Table S1). We analyzed the 3’UTR sequences, confirming the ATGHAH motif associated with the PAS (32).

In addition, as previously observed in other systems (35) a (C/T)(A/G)-rich motif was associated with the maxima of these transcription start site clusters. Overall, 89% of the coding genes have both their TL and 3’UTR sequences supported by identified TSS and PAS clusters, respectively.

The analysis leads to a scheme of a stereotypical *C. neoformans* coding gene (Figure 1A). In average, it is 2,305 bp long (median 2,008 bp) and contains 5.6 short introns (median 5) in its sequence. As previously reported (28), these introns are short (63.4 nt in average) and associated with conserved consensus motifs. The *C. neoformans* TL and 3’UTR have median lengths of 107 nt and 129 nt, respectively (180 nt and 189 nt, mean; Figure 1A,B). Only 887 and 429 genes contain one or more introns in their TL and 3’UTR sequence, respectively; these introns are usually larger (118.3 nt) than those that interrupt the CDS. This gene structure is similar in *C. deneoformans* (Table S1) and there are good correlations between the 3’UTR and TL sizes of the orthologous genes in the two species (Figure 1C,D).

**Figure 1:**
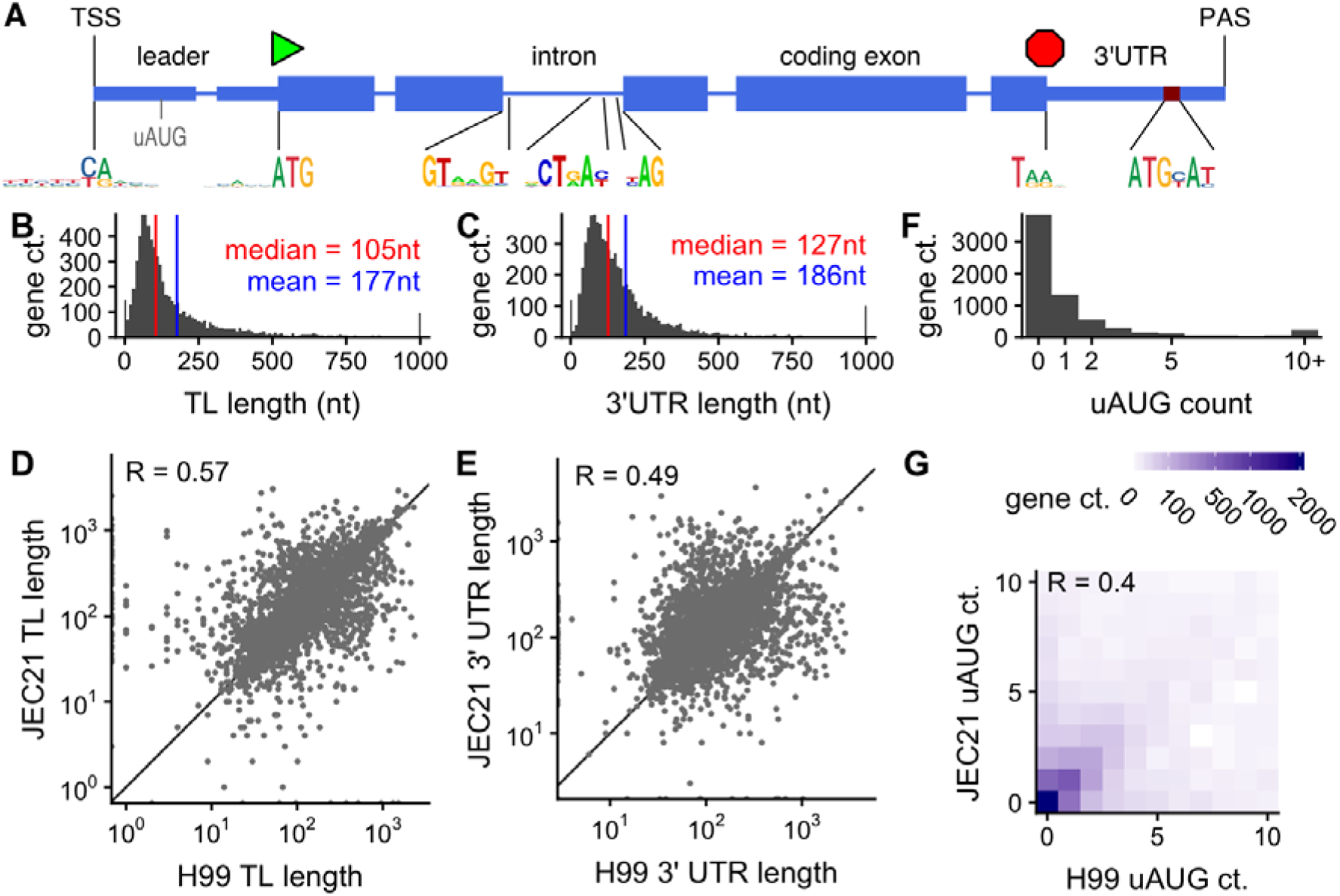
Mapping the coding transcriptome of *Cryptococcus neoformans*. A, Representation of a stereotypical gene of C. *neoformans* H99, showing the sequence logos for the transcription start site (TSS), AUG start codon, intron splicing, stop codon, and polyadenylation site (PAS). B, Distribution of transcript leader (TL) lengths over *C. neoformans* genes, for yeast cells growing exponentially in YPD at 30°C. B, Distribution of 3’ untranslated region (3’UTR) lengths overC. *neoformans* genes. C,D: Comparisons of TLand 3’UTR lengths between orthologous genes in *C. neoformans* H99 and *C. deneoformans* JEC21 growing exponentially in YPD at 30°C. F, Distribution of upstream AUG (uAUG) counts overC. *neoformans* genes, and G, comparison of uAUG counts with *C. deneoformans*.

### More than a third of genes have upstream ORFs that affect translation

The analysis of the TL sequences in *C. neoformans* revealed the presence of 10,286 AUG triplets upstream (uAUG) of the annotated translation start codon (aAUG). We include uAUGs that are either out-of-frame from the start codon, or in-frame but with an intervening stop codon, which are very unlikely to encode a continuous polypeptide. Strikingly, 2,942 genes possess at least one uAUG, representing 43% of the genes with an annotated TL in *C. neoformans* (Figure 1F). A similar result was obtained in *C. deneoformans*, in which we found 10,254 uAUGs in 3,057 genes, and uAUG counts are correlated between the species (Fig 1G).

Translation initiation at uAUGs results in the translation of uORFs, which can regulate expression of the main ORF (36, 37). To evaluate the functionality of the uAUGs in *Cryptococcus, we* generated riboprofiling data in both species and compared densities of ribosome-protected fragments with those of sample-matched poly(A)+ RNA. Our riboprofiling data passes quality metrics of 3-nucleotide periodicity of reads on ORFs indicating active translation by ribosomes, and appropriate read lengths of 26-30nt (Figure S2.1).

Most genes have ribosome occupancy close to that predicted by their RNA abundance, and restricted to the main ORF, for example the most highly translated gene, translation elongation factor eEF1α/CNAG_06125 (Figure 2A,B). However, we observed dramatic examples of translation repression associated with uORFs in CNAG_06246, CNAG_03140 in *C. neoformans* (Figure 2A,C,D). These patterns are conserved in their homologs in *C. deneoformans* (Figure S2.2). Other spectacularly translationally repressed genes, CNAG_07813 and CNAG_07695 and their *C. deneoformans* homologs (Figure 2A, S2.2A) contain conserved uORFs in addition to 5’ introns with alternative splicing or intronically expressed non-coding RNAs (Fig S2.3). In all these cases, high ribosomal occupancy on one or more uORFs is associated with low occupancy of the main ORF.

**Figure 2:**
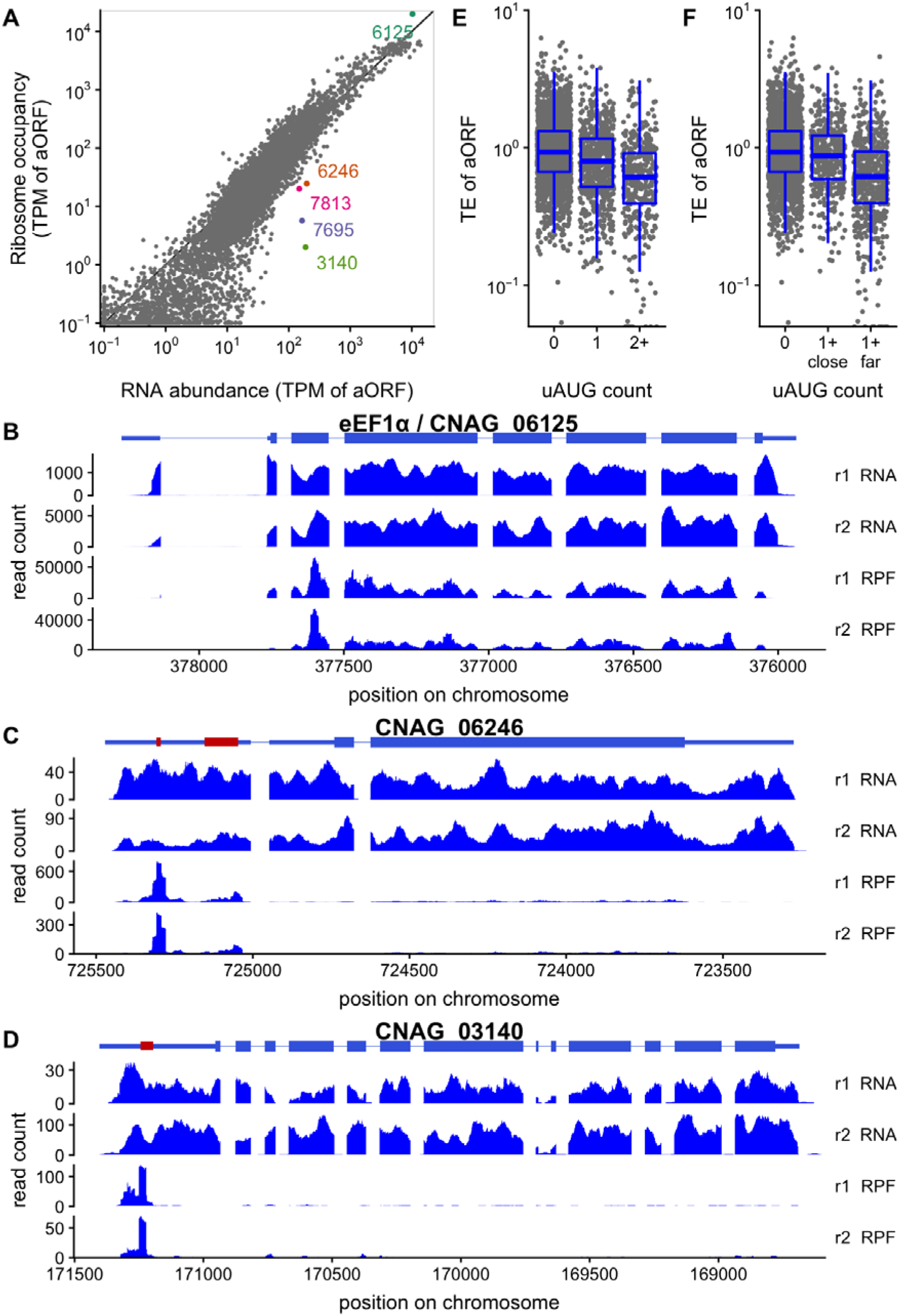
Upstream AUGs repress translation in *C. neoformans*. A, translation regulation of annotated ORFs (aORFs) in *C. neoformans* H99 growing exponentially in YPD at 30°C (equivalent data for *C. deneoformans* shown in Fig S2.1). Ribosome occupancy is plotted against the RNA abundance, both calculated in transcripts per million (TPM) on the aORF. Select genes discussed in the text are highlighted in colour. B, uAUGs are associated with lower translation efficiency (TE) of annotated ORFs, measured as the ratio of ribosome occupancy to RNA-seq reads. C, only uAUGs far from the transcription start site are associated with low TE. A gene is in the “1+ far” category if it has at least one uAUG more than 20nt from the TSS, “1+ close” if all uAUGs are within 20nt of the TSS. D-F, Examples of ribosome occupancy profiles along select RNAs highlighted in 2A (others are shown in Fig S2.2). D, Translation elongation factor eEF2/CNAG_06125 has high ribosome occupancy in the annotated ORF. Translationally repressed mRNAs CNAG_06246 (E) and CNAG_03140 (F) have high ribosome occupancy in uORFs in the transcript leader (red), and low ribosome occupancy in the aORF. Only the first of 5 uORFs in CNAG_03140 is shown. Homologous genes in *C. deneoformans* have similar structure and regulation (Fig S2.1).

The uncharacterized gene CNAG_06246 has two AUG-encoded uORFs that are occupied by ribosomes, and a predicted C-terminal bZIP DNA-binding domain. This gene structure is reminiscent of the multi-uORF-regulated amino-acid responsive transcription factors Gcn4/Atf4 (37), or the S. *pombe* analog Fil1 (38). The sugar transporter homolog CNAG_03140 has six uAUGs, with substantial ribosome occupancy only at the first. Interestingly, *N. crassa* has a sugar transporter in the same major facilitator superfamily regulated by a uORF (rco-3/sor-4, (39)), and sugar-responsive translational repression via uORFs has been observed in plants (40).

Since these translationally repressed genes have multiple uAUGs, we investigated the relationship between uAUGs and translation efficiency genome-wide. We observed a clear negative relationship between the number of uAUGs and translation efficiency (Figure 2E, S2.2E), suggesting an uAUG-associated negative regulation of gene expression in both species.

### Position relative to the TSS affects uAUG translation

Although some uAUGs are recognized and efficiently used as translation start sites, some others are used poorly or not at all, and allow translation of the main ORF. We thus analyzed *Cryptococcus* uAUG position and sequence context to see how translation start codons are specified in these fungi.

We compared the translation efficiency of genes containing only uAUGs close to the TSS to those with uAUGs far from the TSS. In *C. neoformans*, 1,627 of the 10,286 uAUGs are positioned within the first 20 nt of the TL, and 816 uAUG-containing genes have no uAUG after this position. The presence of one or several uAUGs close to the TSS (<20 nt) has nearly no effect on translation efficiency, whereas genes containing uAUGs far from the TSS are less efficiently translated (Figure 2F), and similarly in *C. deneoformans* (Figure S2.2F).

### A Kozak sequence context determines AUG translation initiation

To analyze the importance of AUG sequence context for translation initiation in *Cryptococcus we* used the *5%* most translated genes (hiTrans n = 330) to construct a consensus sequence surrounding their annotated translation start codon (Figure 3A). The context contains a purine at the −3 position, a hallmark of the Kozak consensus sequence (19). However, there is very little enrichment for the +1 position, in contrast with the mammalian Kozak context in which a G is present in +1 ((AZG)C<u>CAUG</u>G) (19). Because of the limited sequence bias downstream of the AUG, and its confounding with signals of N-terminal amino acids and codon usage, we do not consider it further. However, we found a slight sequence bias in the positions −10 to −7 that is outside the metazoan Kozak context. We thus calculated “Kozak scores” for all uAUGs against the position weight matrix (pwm) of the Kozak context from −10 from AUG through to AUG (Figure 3A). We compared the Kozak scores of the annotated AUGs (aAUGs) with those of the 5% most highly translated genes, the first upstream AUG (uAUGs) and the first downstream AUG (d_1_AUG). Highly translated aAUGs have a higher score than typical aAUGs, and aAUGs have usually a higher score than the uAUGs and d_1_AUGs (Figure 3B). This suggests that the sequence context of the AUG codon might be of importance in the selection of the translation start site in *Cryptococcus. We* next asked if the sequence context of uAUGs affected their ability to repress translation of the annotated ORF, focusing on transcripts with only a single uAUG. For uAUGs close to the TSS, there is no correlation between uAUG Kozak score and the translation efficiency of the aORF; there is a weak negative correlation if the uAUG is far from the TSS (Figure 3D). This indicates that the location of an uAUG impacts its activity. The most striking examples of translational repression in Figure 2 tend to have multiple high-score uAUGs (scores CNAG_06246, UjAUG 0.93, u_2_AUG 0.86; CNAG_03140, UjAUG 0.85, u_2_AUG 0.76; CNAG_07813, UjAUG 0.79; CNAG_07695, UjAUG 0.97, u_2_AUG 0.90).

**Figure 3:**
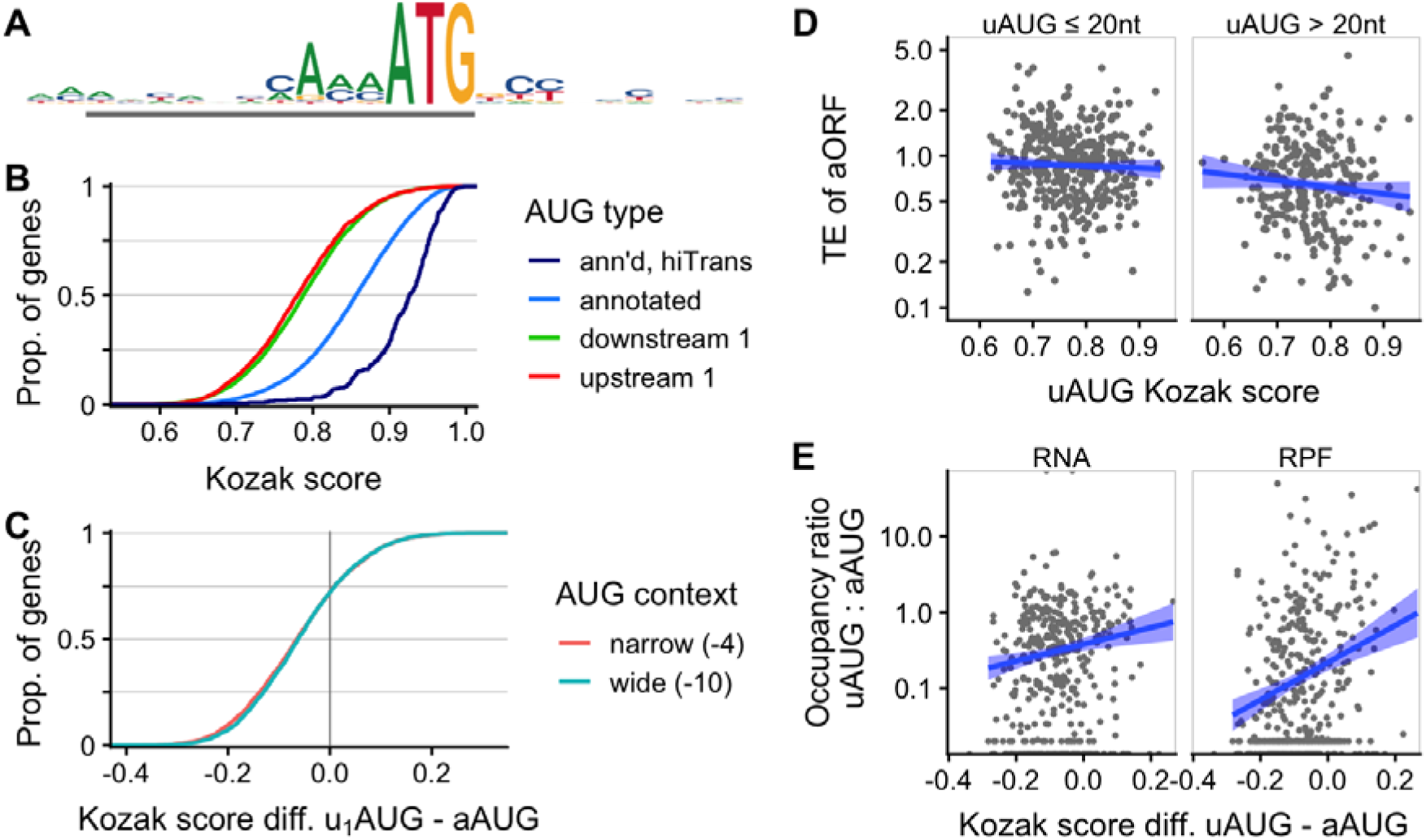
An AUG sequence context is associated with translation in *C. neoformans*. A, Kozak-like sequence context of AUGs, from −12 to +12, for highest-translated 5% of genes (hiTrans). This sequence context is used to create “Kozak scores” of other AUG sequences by their similarity to the consensus from −10 onwards. B, Cumulative density plot of Kozak scores from various categories of AUG, showing that high scores are associated with annotated AUGs of highly translated genes (hiTrans), somewhat with annotated AUGs, and not with the most 5’ downstream AUG (downstream 1) or 5’ most upstream AUG (upstream 1) in a transcript. C, Cumulative density plot of differences in scores between most 5’ upstream (uiAUG) and annotated AUG, showing that for 75% of genes the upstream AUG score is less than the annotated AUG, whether we take a wide (−10:AUG) or a narrow (−4:AUG) window to calculate the score. D, High upstream AUG score is weakly associated with translation repression of the annotated ORF, if the uAUG is further than 20nt from the TSS. E, The relative occupancy of ribosomes (RPF) at the upstream AUG and annotated AUG depends on the difference in scores, even when compared to RNA-seq reads; linear model trend fit shown (blue). Panels D and E show data only for genes in the top 50% by RNA abundance, and with only a single upstream AUG.

We also asked if the AUG score affects the AUG usage transcriptome-wide, by comparing the difference in UiAUG and aAUG scores with the ratio in A-site ribosome occupancy in a 10-codon neighbourhood downstream of the uiAUG and aAUG. We considered the relative occupancy to control for transcript-specific differences in abundance and cap-dependent initiation-complex recruitment. A higher score difference is associated with higher relative ribosome occupancy, while the control comparison with RNA-Seq coverage shows a smaller effect (Figure 3E).

### Nonsense-mediated decay acts on uORF-containing genes

An mRNA molecule translated using an uAUG can be recognized as a premature stop codon bearing molecule and will be as such degraded by the nonsense-mediated mRNA decay (NMD) (41). To test this concept, we first sequenced RNA from *C. deneoformans* strains with the conserved NMD factor Upf1 deleted (33), finding 370 genes with increased mRNA abundance and 270 with decreased (Figure 4A, Table S2; 2-fold difference in levels at 1% FDR).

**Figure 4:**
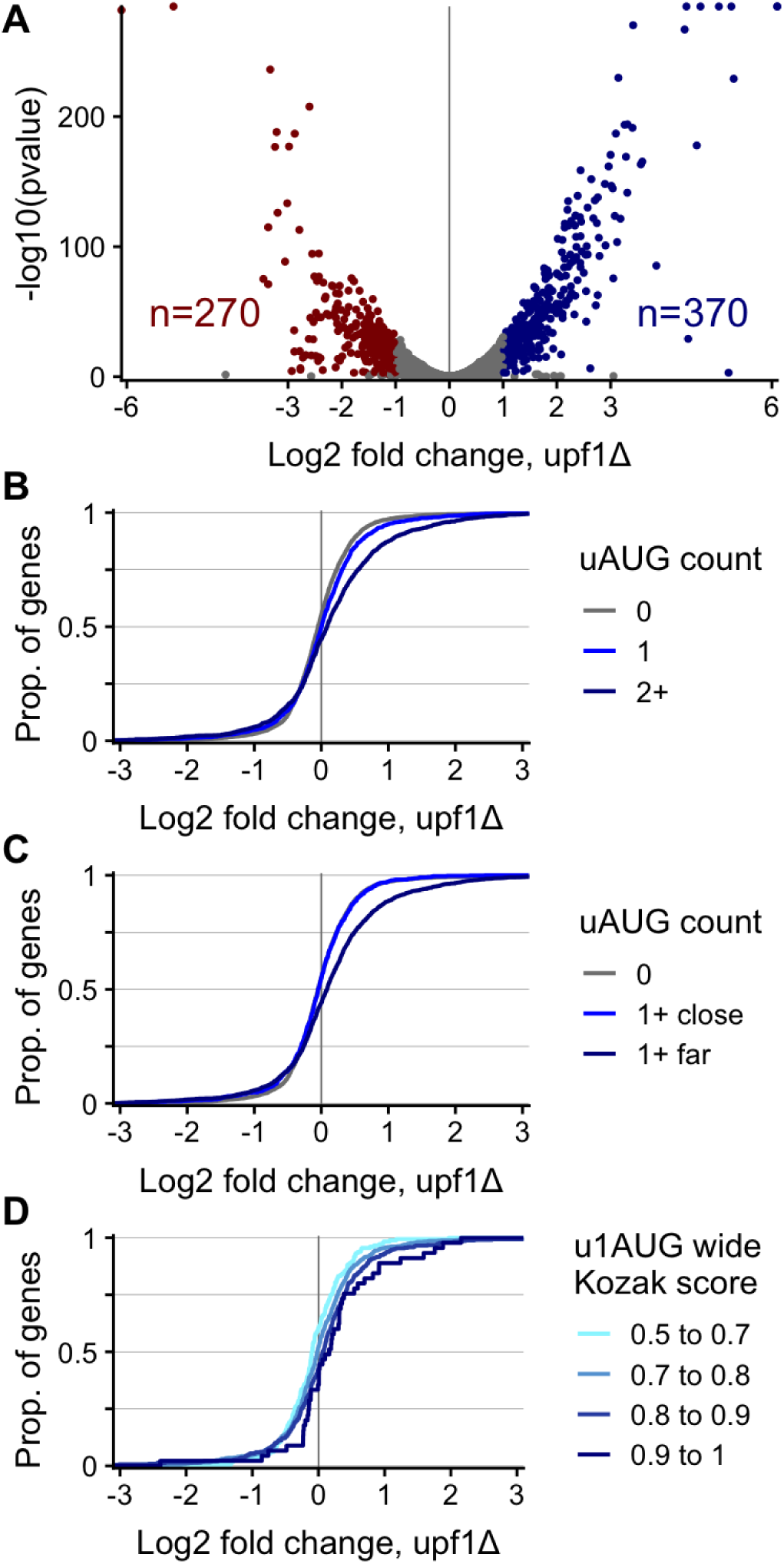
Nonsense-mediated decay (NMD) acts on upstream-ORF-containing mRNAs in *C. deneoformans*. A, Differential expression results from RNA-Seq in *C. deneoformans* JEC21, comparing expression in wild-type cells with a mutant deleted for NMD factor DPF1/CNC02960, and using DeSeq2 to identify genes upregulated in *the upf1Δ* mutant. B, uORF containing genes are enriched for NMD-sensitivity. C, uORF-containing genes are enriched for NMD-sensitivity only when the uAUG is more than 20nts from the TSS (1+far), but not when the uAUG is less than 20nts (1+close). D, Start codon sequence context affects NMD sensitivity of genes containing a single upstream AUG: uORFs starting with higher Kozak-score uAUG are more likely to increase in abundance in the *upf1Δ* mutant.

We next compared the fold-change in abundance of uORF-containing or uORF-free mRNAs. Two genes with extreme increases in *upflΚ* are also extremely translationally repressed uORF-containing genes we identified above (Figures 2, S2.1, S2.2): CNF00330 (CNAGJD7695 homolog, 11-fold) and CNG04240 (CNAGJD3140 homolog, 8-fold). Another extreme is the carbamoyl-phosphate synthase CND01051 (5-fold up in *upf1Δ)*, a homolog of S. *cerevisiae CPA1* and *N. crassa arg-2*. These orthologs are regulated by a conserved uORF encoding a arginine attenuator peptide that have all been verified to repress reporter gene synthesis in a *N. crassa* cell-free translation system (42). Consistent with this model, in both *C. neoformans* and *C. deneoformans* the native uORF shows strong ribosome occupancy while the aORF is translationally repressed (CnTE = 0.47, CdTE = 0.38; Figure S4.1).

In general, uORF-containing genes are more likely to be up regulated in the *upf1Δ* mutant than uAUG-free genes (Figure 4B), suggesting that uORFs negatively regulate mRNA abundance in *Cryptococcus*, in addition to repressing translation of the main ORF. Similarly, uORF-containing genes are enriched for NMD-sensitivity only when the uAUG is more than 20 nt from the TSS (Figure 4C), suggesting that TSS-proximal uAUGs (< 20 nt) are skipped, and generally not used as translation start codons in *Cryptococcus*.

Next, we asked if uAUG Kozak score affects mRNA decay via the NMD pathway. Restricting our analysis to genes with a single uAUG (n=1,421), we binned genes according to their Kozak score. We find that mRNAs that contain uORFs which start with a higher Kozak-score uAUG are more likely to increase in abundance in the *upf1Δ* mutant (Figure 4D). Indeed, the abundance increase is monotonically correlated with the mean of the score bins. In conclusion, in *Cryptococcus*, the position and the sequence context of uAUGs determines their usage as translation start codons, and the effect of the uORF on stimulating nonsense-mediated decay of the mRNA.

### Start codon sequence context and uORF regulation in other fungi

We then examined sequences associated with translation start codons in other fungi, for which both RNA-Seq and Riboprofiling data were available, and for which the annotation was sufficiently complete (i.e. S. *cerevisiae; Neurospora crassa, Candida albicans* and *Schizosacchomyces pombe). We* analyzed the Kozak context associated with aAUG of all annotated coding genes, of the *5%* most translated genes (hiTrans), and for mRNAs encoding cytoplasmic ribosomal proteins (CytoRibo), as a model group of highly expressed and co-regulated genes defined by homology (Table S3). Cytoplasmic ribosomal proteins have informative Kozak contexts, with a strong A-enrichment at the positions −1 to-3 and weak sequence enrichment after the AUG in all these species (Figure 5A). The total information content of the Kozak sequence is higher for CytoRibo genes than HiTrans, and higher for HiTrans than all annotated genes, in all these fungi (Figure 5B). Nevertheless, these contexts have also some species specificity: Kozak sequences for HiTrans and CytoRibo are more informative in *Cryptococcus* and *N. crassa* than in 5. *pombe, C. albicans*, and 5. *cerevisiae*. In particular, the C-enrichment at positions −1, −2 and −5 in *Cryptococcus* and *N. crassa* is absent in 5. *cerevisiae*, and we observed no sequence enrichment upstream of the - 4 position forS. *pombe* and very little forS. *cerevisiae*. In contrast, a −8 C enrichment, similar to the *Cryptococcus* pattern, was observed in *N. crassa*. The −10 −6 A/T rich region for *C. albicans* is likely to reflect an overall A/T-richness of the TLs in *C. albicans*.

**Figure 5:**
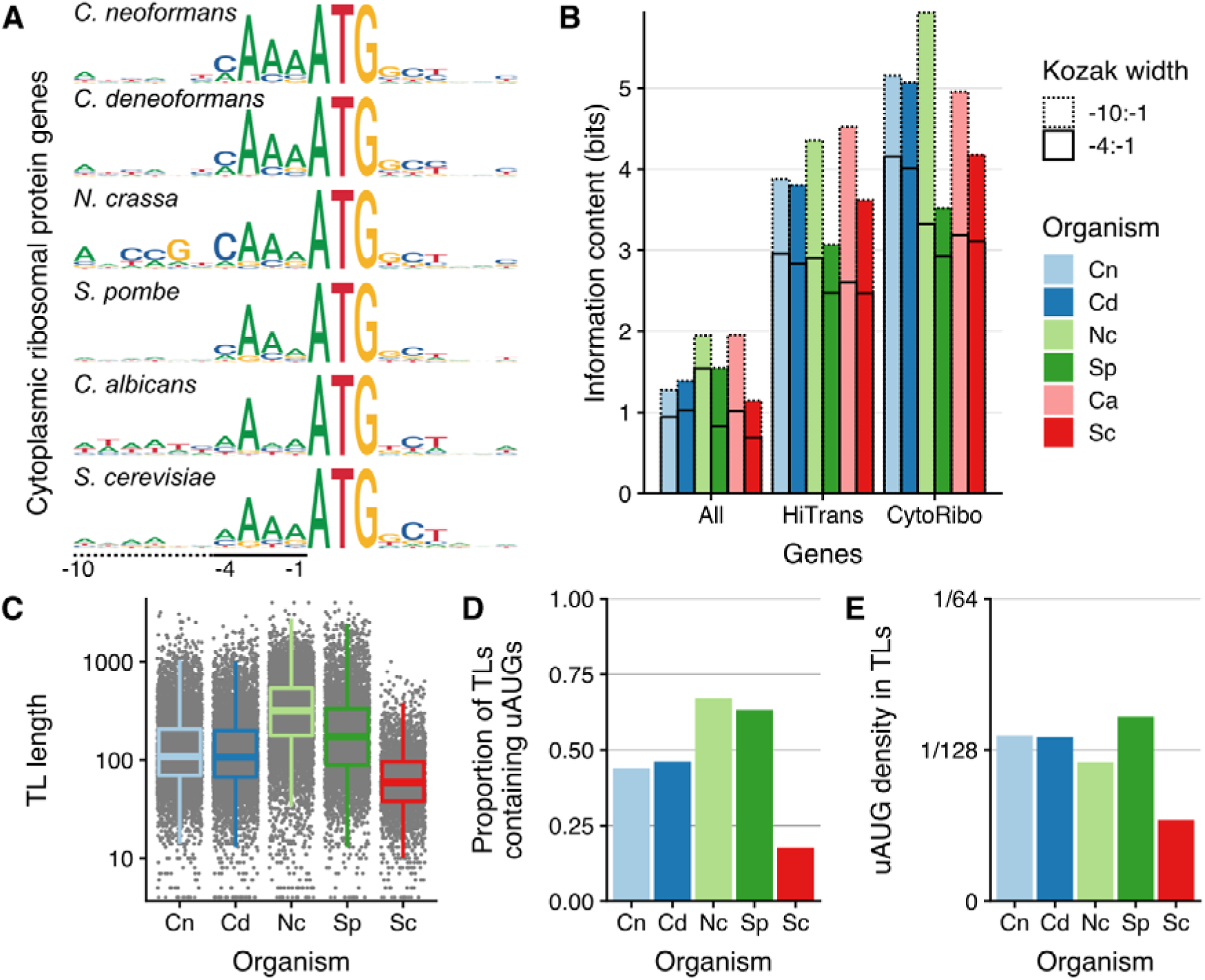
Sequences specifying start codon selection are quantitatively different in different fungi. A, Kozak consensus sequence logo for annotated start codons of cytoplasmic ribosomal protein genes from 6 fungal species. The height of each letter represents the Shannon information content in bits, so that the anchor ATG sequence has height 2 bits. B, Information content at annotated start codons in bits per base (i.e. summed height of stacked letters in sequence logo) for 3 groups of genes, in the 6 fungi from panel A. Solid line indicates information from −1 to −4 of ATG, and dotted line additionally to −10 (see bottom of panel A). Gene groups are all annotated ORFs, highly translated ORFs (HiTrans) and cytoplasmic ribosomal proteins (CytoRibo, as panel A). HiTrans used the highest-translated 5% of genes, or the highest 400 genes for fungi with more than 8000 annotated genes (C. *albicans* and *N. crassa;* see methods). C-E, For 5 fungi for which transcript leader (TL) annotations were available, TL length (C), proportion of annotated TL containing an upstream AUG (D), and proportions of AUGs per nucleotide in the TL (E; a uniform random model would have density 1/64).

The analysis of the TL sequences from these fungi, excluding *C. albicans* for which no TL annotation is available, also shows species specificity. The average TL length in S. *cerevisiae* (84 nt) is less than half that in *Cryptococcus* (Figure 5C). In sharp contrast with *Cryptococcus*, only 985 uAUGs are present in 504 genes, which correspond to 18% of the genes with an annotated TL in 5. *cerevisiae*. Moreover, the density of the uAUGs is very low and uAUGs have no global effect on TE in this yeast (Figures 5D,E, Fig S5.1).

More broadly, short TLs with very low uAUG density are more the exception than the rule in the fungal kingdom (Figure 5C). Most fungi have large number of uAUGs in their TL and these uAUGs globally down regulate gene expression (Figure S5.1). Our analysis in *Cryptococcus*, together with the strength and quality of the Kozak contexts associated with the aAUG, suggest that fungi in general are able to discriminate between AUGs with different sequence context, and thus are using AUG sequence context to regulate gene expression at the post transcriptional level.

### Kozak context controls leaky scanning in *Cryptococcus*

We earlier calculated the Kozak score of the first downstream AUG (d_1_AUG) within each CDS: these d_1_AUG scores are mostly lower than the score of the aAUGs (Figure 3B), consistent with most annotations correctly identifying a good-context AUG as the start codon. Yet, we identified number of d_1_AUGs with a high score (n=1109 above 0.826, the median Kozak score for aAUGs; n= 131 above 0.926, the median for hiTrans), which could be efficiently used as translation start codon. The scanning model of translation initiation predicts that the d_1_AUG will be used as the start codon only if pre-initiation complexes leak past the aAUG. Thus if the aAUG has a strong sequence context, most ribosomes will start translation there and the d_1_AUG will be not used as a translation start codon.

To identify potential leaky translation initiation events, we compared the aAUG and d_1_AUG scores for each of the 50% most expressed genes (Figure 6A). For above-median aAUG score genes, the score of the d_1_AUGs can be very high or very low. By contrast, for the genes with a low aAUG score, there is a bias toward higher d_1_AUG score, suggesting that for these genes the strong d_1_AUG could be used as alternative translation start site (Figure 6A).

**Figure 6:**
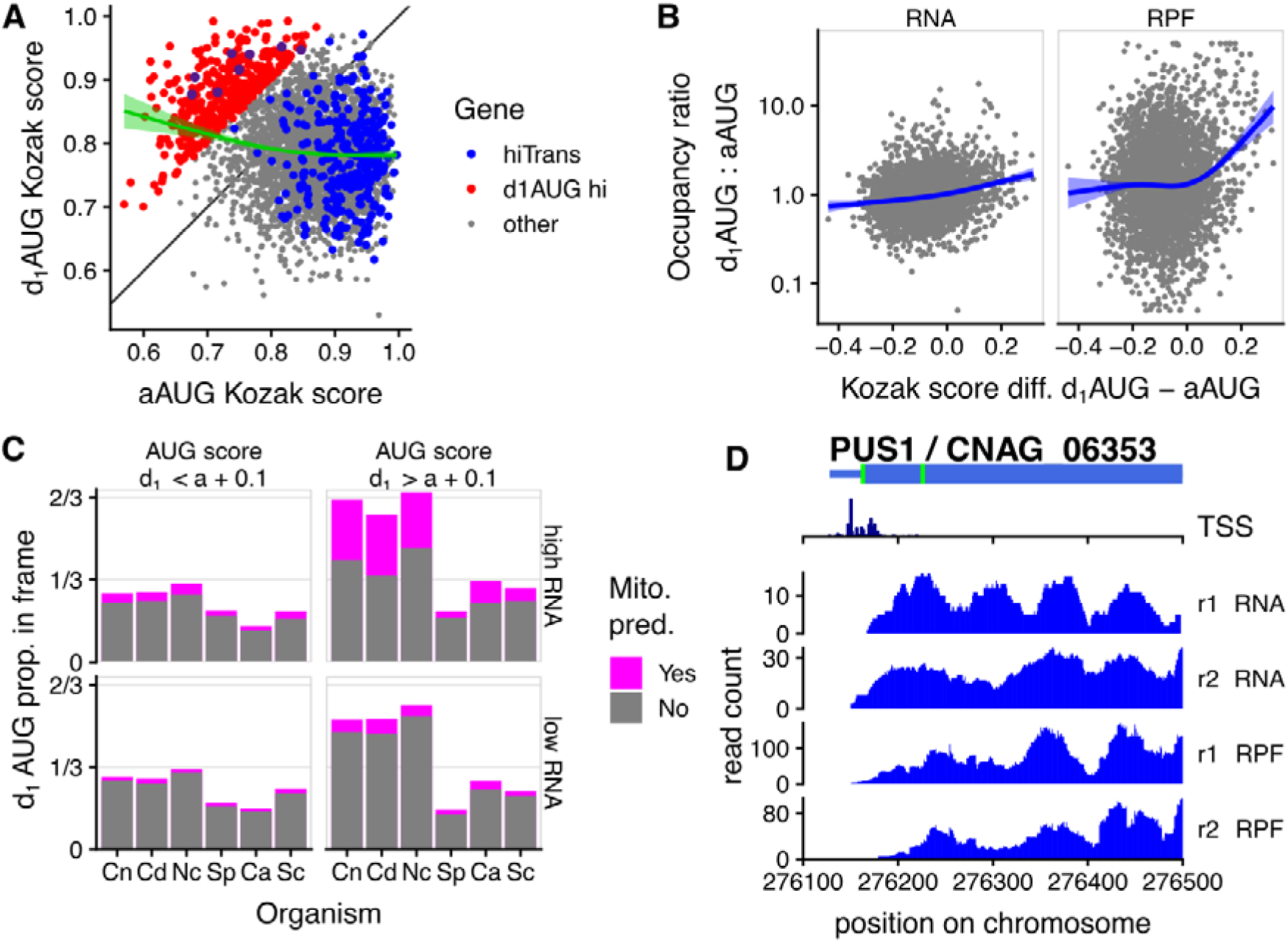
High-scoring downstream AUGs specify alternative N-terminal isoforms in *C. neoformans*. A, Most genes with reasonable RNA abundance (top 50% by RNA abundance shown), especially very highly-translated genes (blue, top 5%), have lower Kozak score at the 1st downstream AUG than at the annotated AUG. However there are exceptions (red, d_1_AUG hi: d_1_AUG score > annotated AUG score + 0.1), and there is a trend for genes with low aAUG score to have a higher d_1_AUG score (green, generalized additive model fit). B, Higher d_1_AUG score than aAUG score drives higher ribosome protected fragment (RPF) occupancy at the d_1_AUG compared to the aAUG, but much smaller differences in RNA-seq density. Blue line indicates generalized additive model fit. C, Downstream AUGs with high Kozak scores (d_1_AUG score > annotated AUG score + 0.1) and reasonable RNA abundance (top 50%) are likely to be in-frame and enriched for N-terminal mitochondrial localization signals in *C. neoformans, C. deneoformans*, and *N. crassa*, but not in 5. *pombe, C. albicans*, or 5. *cerevisiae*. D, The pseudouridine synthase CnPusl is a candidate alternate-localized protein with a low-score aAUG and high-score d_1_AUG, and transcription start sites on both sides of the aAUG. RNA-Seq and RPF reads on the first exon are shown, and the full length of the gene shown in Fig S6.1.

To test whether AUG score affects translation initiation, we calculated the ratio of ribosome protected fragment density and RNA-Seq density around each aAUG and d_1_AUG, and the difference in score between these two AUGs (Figure 6B). We found a weak positive correlation between the difference in scores of the two AUGs and RNA-Seq density at these specific loci, raising the possibility that transcription start sites sometimes occur downstream of a weak aAUG. By contrast, the relative ribosome density sharply increases at the d_1_AUGs when the difference in score between the d_1_AUG and the aAUGs increases. This suggests that for these genes, both AUGs can be used as translation start codon, as because a subset of scanning ribosomes leak past the aAUG and initiate at the d_1_AUG.

### Kozak context-controlled scanning specifies alternative N-termini in *Cryptococcus* and ***Neurospora***

*We* next determined which groups of genes could be affected by potential alternative start codon usage. We focused our analysis on the 50% most expressed genes for which the difference in score between the aAUG and d_1_AUG was the highest (difference in wide score d_1_AUG - aAUG > 0.1, n = 167 for *C. neoformans)* (Table S4). Strikingly, for 66% of these genes (110/167) the d_1_AUG is in frame with the corresponding aAUG, with a median of 69 nt (mean 79 nt) between the two AUGs. Thus, alternative usage of in-frame AUGs would result in proteins with different N-terminal ends. Supporting this hypothesis, 37% of these proteins (41/110) possess a predicted mitochondrial targeting sequence located between the two AUGs, far exceeding the 8% genome-wide (560/6788). This suggests that the usage of the annotated start codon would target the isoform to mitochondria, whereas the usage of the d_1_AUG would produce a protein specific to the cytoplasm or another organelle. Examples of alternative localization driven by alternative N-termini have been observed across eukaryotes (43).

The pattern of predicted dual-localization, i.e. enrichment of high-score d_1_AUGs in-frame with predicted mitochondrial localization signal on the longer N-terminal, is conserved in some fungi but not others (Figure 6C). In a null model where coding sequences have random nucleotide content, we would expect roughly 1/3 of d_1_AUGs to be in frame. In 6 fungal species we examined, for d_1_AUGs whose score is comparable to or less than the aAUG they follow, the proportion in frame is close to *(Cryptococcus, N. crassa)* or less than 1/3. These proportions are similar when we considered reasonably expressed (above-median) or low expressed genes. The pattern differs for proteins with a d_1_AUGs whose score high relative to the aAUG they follow (d_1_AUG score > aAUG score + 0.1). In *Cryptococcus* and *N. crassa*, most reasonably expressed mRNAs are in-frame and over 1/3 of these in-frame high-score d_1_AUGs have predicted mitochondrial localization. InS. *cerevisiae* and *C. albicans*, a relative slight enrichment for in-frame d_1_AUGs and for protein possessing a mitochondrial targeting sequence for high-scoring d_1_AUGs can be also observed. By contrast, in S. *pombe we see* depletion in the in-frame/out-of-frame ratio, even in these proteins with high-scoring d_1_AUGs.

These results suggest that the extent to which alternate translation start codons regulate proteome diversity is variable in fungi. Accordingly, we identified a number of *Cryptococcus* proteins with potential alternative start codons and N-terminal targeting sequences, whose two homologs in S. *cerevisiae* are known to be necessary in two compartments of the cells. For instance, CnPL/51/CNAG_06353 is an homolog of both the mitochondrial and cytoplasmic tRNA:pseudouridine synthases encoded by the *PUS1* and *PUS2* paralogs in S. *cerevisiae*. In *C. neoformans*, ribosome occupancy at both the aAUG and d_1_AUG of CN AG_06353, and the presence of transcription start sites both sides of the aAUG (Figure 6D), argues that both AUGs are used as start codons, and transcription and translation regulation could co-operate to set isoform levels. Similarly, CnGI01/CNAG_04219 encodes both the cytoplasmic and nuclear isoforms of the glyoxalase I depending of the alternate AUG usage (Figure S6.1B). The next enzyme in this pathway, Glyoxalase II, is likewise encoded by CnGI02/CNAG_01128, which is a homolog of both cytoplasmic (Glo2) and mitochondrial (Glo4) enzymes in S. *cerevisiae*. CNAG_01128 has a weak aAUG, strong d_1_AUG, and N-terminal predicted mitochondrial targeting sequence (Figure S6.1C). Finally, we observed that nine members of the amino-acyl tRNA synthetase gene family have predicted alternate localization from alternate AUG start codons.

### Amino-acyl tRNA synthetases (aaRSs) are frequently single-copy and dual-localized in ***Cryptococcus***

The tRNA charging activity of aaRSs is essential in both cytosol and mitochondria to support translation in each compartment, and examples of alternative localization of two aaRS isoforms of a single gene have been observed in fungi, plants, and animals (44–46). This implies that a eukaryote with a single genomic homolog of an aaRS is likely to make distinct localized isoforms from that locus. Thus, we examined predicted aaRS localization in fungi. We assembled gene lists of aaRSs in diverse fungi from homology databases OrthoDB (47) and PANTHERdb (48), adding a mitochondrial SerRS (CNAG_06763/CNB00380) to the list of *Cryptococcus* aaRSs analysed by Datt and Sharma (49).

In *C. neoformans* and *C. deneoformans*, eleven aaRSs are each expressed from a single genomic locus, including the homologs of all fiveS. *cerevisiae* aaRSs whose dual-localization has been verified (Table S5). Nine of these *Cryptococcus* aaRSs have the same structure of a poor-context annotated AUG followed by a predicted mitochondrial targeting sequence and a strong-context d_1_AUG (Figure 7A/B; AlaRS, CysRS, GlyRS, HisRS, ValRS, LysRS, ProRS, ThrRS, TrpRS). The similar annotated AUG contexts, sharing an unfavourable −3U, suggests that the same mechanism could lead to leaky translation initiation at most of these (Table S6). At the downstream AUGs, the strong Kozak context is consistent with efficient translation initiation of the cytoplasmic isoform from this start codon (Table S6).

**Figure 7:**
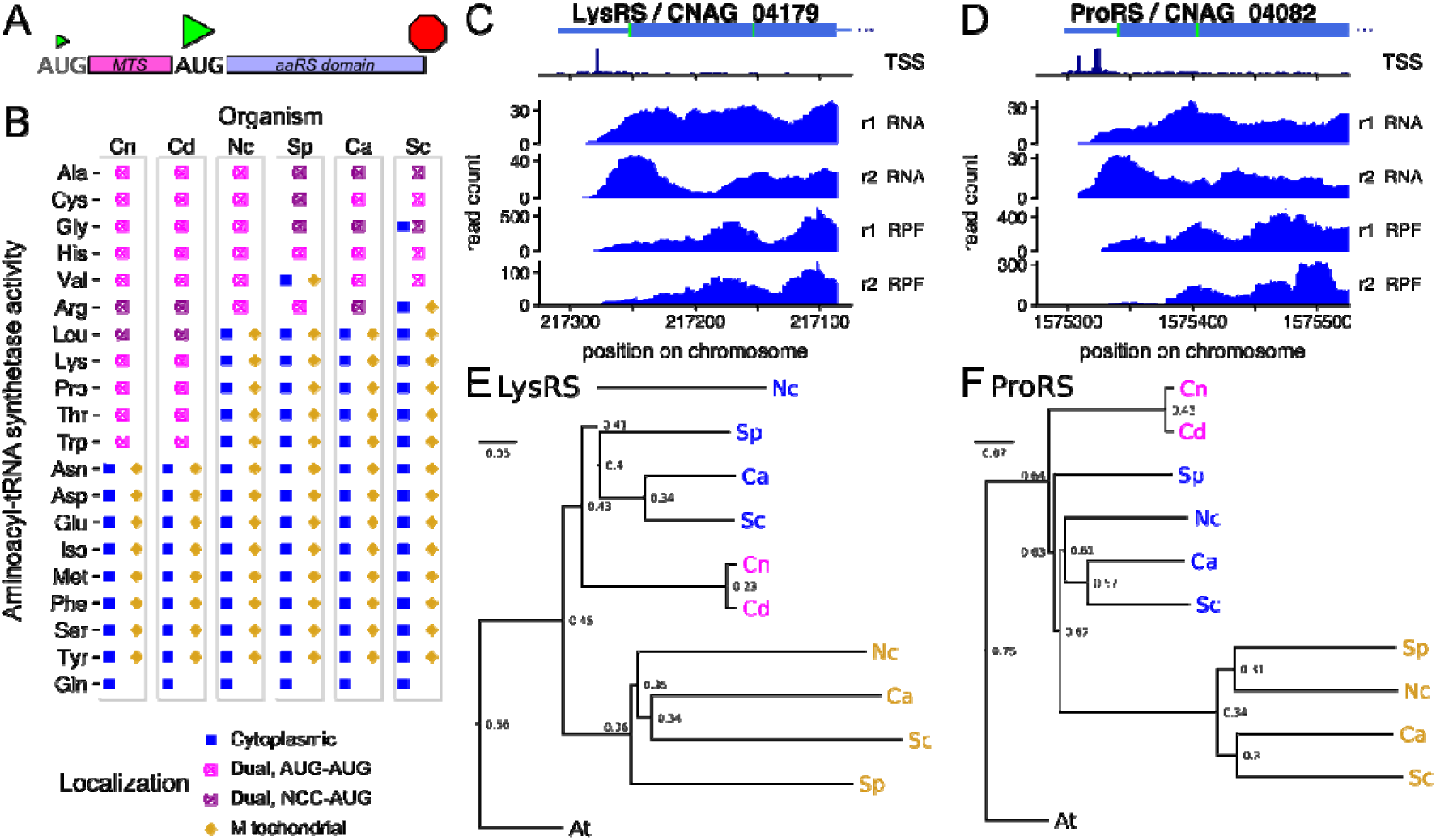
Aminoacyl-tRNA synthetases (aaRSs) are commonly alternatively localized to cytoplasm and mitochondria by use of alternative start codons in fungi. A, Schematic of the structure of a dual-localized aaRS with alternate AUG start codons. B, Predicted localization of all aaRS enzymes in the fungi *C. neoformans* (Cn), *C. deneoformans* (Cd), *N. crassa* (Nc), S. *pombe* (Sp), *C. albicans* (Ca), S. *cerevisiae* (Sc). C/D, Transcription start site reads, RNA-seq, and ribosome profiles of 5’-ends of CnLysRS (C) and CnProRS (D) show that most transcription starts upstream of both AUG start codons (green), and both AUG codons are used for translation initiation. E/F Simplified neighbour-joining phylogenetic trees show that LysRS (E) and ProRS (F) genes were duplicated in ascomycete fungi, and *Cryptococcus* retained a single dual-localized homolog. *Arabidopsis thaliana* (At) was used as an out-group. The scale bar represents the number of amino acid substitutions per residue, and the numbers at nodes are the proportion of substitutions between that node and its parent. See table S5, for details of identifiers for genes (GenelD).

The two remaining single-copy aaRSs have near-AUG translation initiation sites upstream of predicted mitochondrial targeting sequences. Translation of ArgRS starts at an AUU codon with otherwise strong context (cccaccAUU) conserved in both *Cryptococcus* species. This N-terminal extension includes a predicted mitochondrial targeting sequence (mitofates p > 0.95 for both species). Translation of LeuRS starts at adjacent ACG and AUU codons which collectively provide strong initiation context (gccaccACGAUU in *C. neoformans*, gccACGAUU in *C. deneoformans)*. This N-terminal extension also includes a predicted mitochondrial targeting sequence (mitofates p≈0.7 for both species).

In *Cryptococcus*, alternative aaRS isoforms appear to be mostly generated by alternative translation from a single transcript, and sometimes by alternative transcription start sites. On all the predicted dual-localized aaRSs, we observe ribosomal occupancy starting at the earliest start codon (Figure 7C/D, Fig S7.1). LysRS/CNAG_04179 contains only a single cluster of transcription start sites, upstream of the aAUG (Figure 7C). ProRS/CN AG_04082 contains a wider bimodal cluster of TSSs, both upstream of the aAUG. Similarly, most transcription initiation is well upstream of the aAUG in CysRS/CNAG_06713, LeuRS/CNAG_06123, ThrRS/CNAG_06755, and ValRS/CNAG_07473. However, for GlyRS/CNAG_05900, and HisRS/CNAG_01544, we observe alternative transcription start sites closely upstream of the annotated start codon, that are likely to affect the efficiency of start codon usage. In ArgRS/CN AG_03457 there is also an alternative transcription start site, close to the near-AUG start codon for the mitochondrial form. In AlaRS/CNAG_05722 and TrpRS/CNAG_04604 we detect some transcription start sites between the alternative start codons, and TrpRS also has an uORF in the transcript leader that is likely to affect translation. These observations suggest that dual-localization of the single-copy aaRSs in *Cryptococcus* is regulated largely by start codon choice. For some genes this regulation is backed up by alternative TSS usage.

Some dual-localized genes use an upstream near cognate codon (DualNCC) in all these fungi, but the NCC-initiated aaRS are not the same from one fungus to the other. For instance, both *Cryptococcus* and *N. crassa* AlaRS use DualAUG whereas in S. *pombe, 5. cerevisiae* and *C. albicans* a DualNCC is used. On the other hand, S. *pombe* GlyRS is regulated by DualNCC whereas the other ones use a DualAUG regulation. Substitution between weak AUG codons and near-cognate codons seems thus to have taken place multiple times in the fungal kingdom.

### Amino-acyl tRNA synthetases as an evolutionary case study

To understand patterns of dual-localization, we next examined the evolution of aaRSs. The ancestral eukaryote is thought to have had two complete sets of aaRS, one mitochondrial and one cytoplasmic, but all mitochondrial aaRSs have been captured by the nuclear genome and many have been lost (50). Thus we examined aaRS phylogenetic trees in more detail. For some amino acids (Asn, Asp, Glu, Iso, Met, Phe, Ser, Tyr), reference fungi have distinct cytoplasmic and mitochondrial aaRSs that cluster in separate trees (51). We also do not consider Gln, because organellar Gln-tRNA charging in some species is achieved by an indirect pathway (52).

Dual-localized AlaRS, CysRS, and HisRS in the 6 fungi we focus on are each monophyletic (51). Even these aaRS can be encoded by two genes in some other fungi: AlaRS is duplicated to one exclusively mitochondrial and another exclusively cytoplasmic gene in the Saccharomycete yeast *Vanderwaltozyma polyspora* (53). For CysRS, *Aspergillus versicolor* (ASPVEDRAFT_141527 and ASPVEDRAFT_46520) and *Coprinus cinerea* (CC1G_O3242 and CC1G_14214) have two copies, one of which has a predicted mitochondrial targeting sequence. For HisRS, *Rhizopus delemar* (R03G_01784 and RO3G_16958) and *Phycomyces blakesleeanus* (PHYBL_135135 and PHYBL_138952) likewise contain gene duplications. Similarly, S. *cerevisiae* has two ArgRS genes that arose from the whole-genome duplication: RRS1/YDR341C is essential, abundant, and inferred to be cytoplasmic (54) while MSK1/YHR091C has a mitochondrial localization sequence and MSR1 deletions have a petite phenotype (55), although both have been detected in mitochondria suggesting some residual dual-localization of the cytoplasmic enzyme (56). The second S. *cerevisiae* stress-responsive cytoplasmic copy of GlyRS also arose from the whole-genome duplication (57).S. *pombe* cytoplasmic ValRS is monophyletic with dual-localized ValRS in other fungi, and *Schizosaccharomyces* also has a paralogous but diverged mitochondrial ValRS that appears to be descended from an early eukaryotic ValRS of mitochondrial origin (58).

LysRS appears to have been duplicated in an ancestor of ascomycetes: ascomycete mitochondrial homologs cluster together, and ascomycete cytoplasmic homologs cluster together, while the single basidiomycete homolog clusters close to the base of this split from other opisthokonts (51). By contrast, LeuRS, ProRS, and TrpRS are each represented by two distinct proteins in ascomycetes, one cytoplasmic and one mitochondrial and of independent descent, but the mitochondrial homolog has been lost in *Cryptococcus* species. In basidiomycetes *Ustilago* and *Puccinia*, homologs of mitochondrial LeuRS and ProRS are not present, but there is a homolog of mitochondrial TrpRS; all these have a single homolog of the cytoplasmic TrpRS (51). Our independent phylogenetic analysis of LysRS and ProRS agrees with the conclusions from PANTHERdb (Figures 7E/F). These analyses show that aaRSs have undergone multiple incidences of at least two processes during fungal evolution: losses associated with the dual-localization of the remaining gene, and duplications followed by specialization.

### Evolutionary conservation of gene-specific feedback regulation by alternate AUG usage

We also observed striking examples of gene-specific regulation by start codon context in *Cryptococcus*, in translation factors affecting start codon selection, supporting previously proposed models of feedback regulation (59, 60).

Translation initiation factor eIF1, which enforces selection of strong context start codons, is encoded by an mRNA with poor start codon context in diverse eukaryotes, driving an autoregulatory program (59, 61). In *C. neoformans*, eIF1 also initiates from a poor-context cuuaguugaAUG start (score 0.75), and ribosome profiling reads are spread across the annotated ORF (Figure 8A). I nt rigui ngiy, the next AUG is out-of frame and has strong context cuccaaaaAUG (score 0.98), with a same-frame stop codon 35 codons later, suggesting that this could represent a downstream short ORF that captures ribosomes that have leaked past the poor-context start. To test this hypothesis, we examined the 5’ ends of riboprofiling reads, which report on the translation frame of the ribosomes (36). Riboprofiling reads from the 5’ and 3’ of the eIF1 annotated ORF are roughly 77% in frame 0,10% in +1, and 13% in +2, as are reads on two other highly expressed genes, eEF1α and *HSP90*. By contrast, in the hypothesized downstream ORF, reads are only 57% in frame 0, 32% in frame +1, and 11% in frame +2, consistent with translation occurring in both frame 0 and +1. The gene structure is conserved in *C. deneoformans*, with a weak aAUG (score 0.76), a strong d_1_AUG (score 0.98) in the +1 frame, followed by an enrichment in +l-frame riboprofiling reads (Figure S8.1A,B). We observe small increases in eIF1 mRNA levels in the *upf1Δ* strain of *C. deneoformans* at both 30°C (1.16x, p=0.04) and 37°C (1.09x), so NMD could regulate this transcript. Overall, our data support the hypothesis that the downstream ORF of eIF1 is translated after read-through of the annotated AUG, and that the downstream ORF contributes to translation regulation of the annotated ORF.

**Figure 8.**
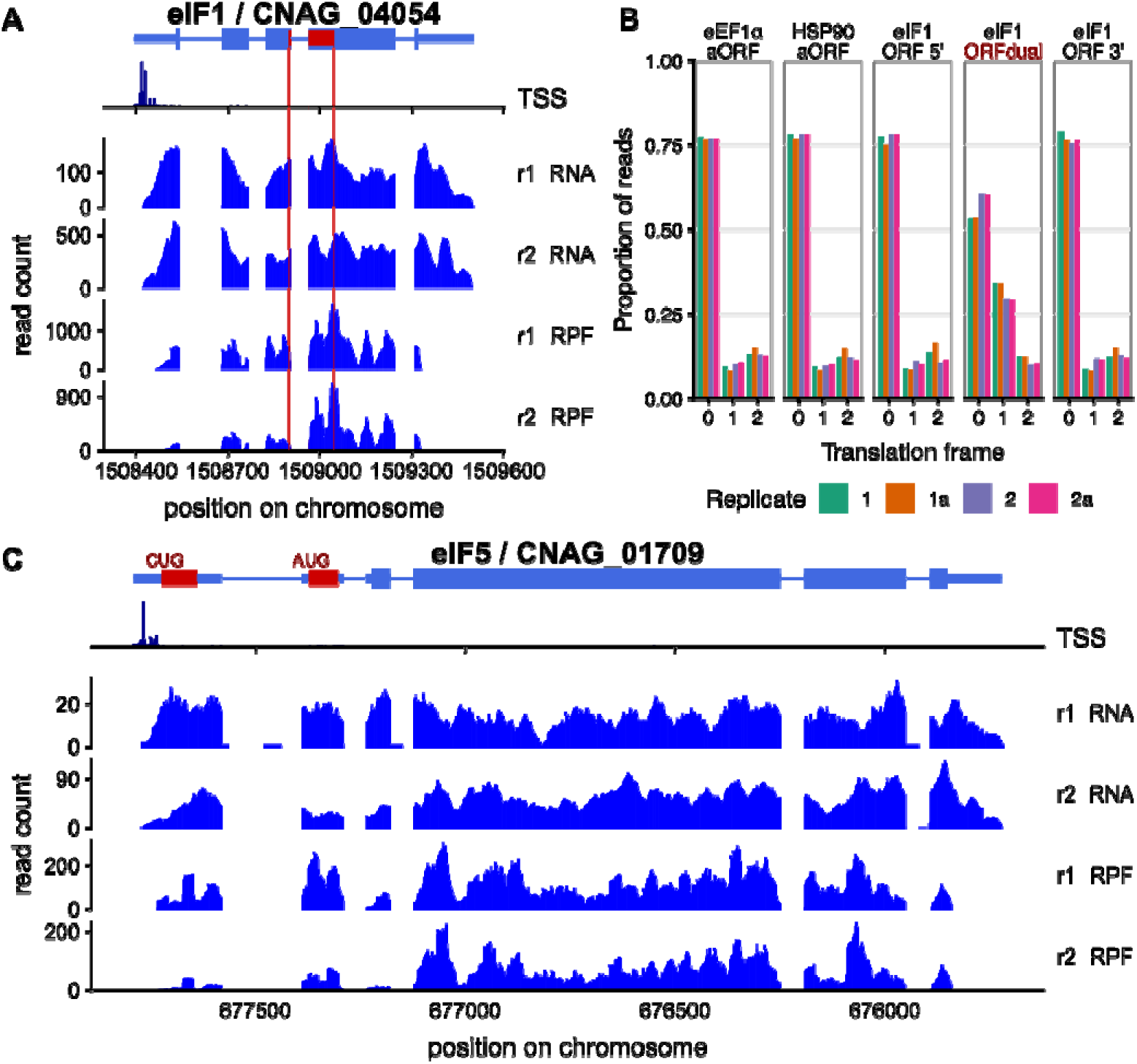
Translation initiation factors eIF1 and eIF5 are regulated by alternate start codon usage in *C. neoformans*. A, Reads on CneIF1/CNAG_04054, showing frame +1 “downstream ORF” in dark red, breaking for an intron. B, The downstream ORF of CneIF1 is dual-translated in two frames. Most ribosome profiling read 5’ ends are in a consistent frame, including in control genes eEF1alpha/CNAG_06125 and HSP90/CNAG_06125, and in the 5’ and 3’ ends of the CneIF1 ORF, but there is 2x enrichment of reads in frame+1 in the dual-decoded ORF. C, Reads on CneIF5/CNAG_01709 showing substantial ribosomal occupancy over upstream ORFs. The first upstream ORF shown is translated from a CUG start codon and the second from an AUG codon, and other uORFS potentially initiated from near-cognate codons are not shown. C. *deneoformans* homologs have the same structure and regulation (Figure S8).

Translation initiation factor eIF5 reduces the stringency of strong-context start-codon selection, and is encoded by an mRNA with a repressive uORF initiated from a poor-context uAUG in diverse eukaryotes (60). In *Cryptococcus*, eIF5 (77F5/CNAG_01709) also contains a uAUG with the poor sequence context aaagaguucAUG (score 0.72), while the main ORF of eIF5 is initiated by a strong context cccgcaaaAUG (score 0.94). We detect ribosomal density on the uORF of *TIF5* comparable to that on the main ORF (Figure 8C), suggesting there is substantial translation initiation at the uAUG. There is also clear translation initiation at a further upstream near-cognate CUG codon. The gene structure is conserved in *C. deneoformans TIF5*, with the same pattern of riboprofiles at upstream poor-context AUG and near-cognate codons (Figure S8). Further, the *C. deneoformans* homolog transcript abundance increases substantially in the *upf1Δ* strain (2.6x, p < 10^−50^). This supports the model that eIF5 translation is repressed by upstream reading frames initiated from poor start codons, leading to nonsense-mediated decay of the transcript.

These examples further illustrate that the first start codon is not always used, but rather start codon usage is driven by the sequence context, and that variability in start codon context including the canonical AUG sequence is used for translational regulation.

### Variable inserts in eTIFs correlate with variation in translation initiation determinants

The conserved proteins eIF1, eIF5, and eIF1A play pivotal roles in start codon selection inS. *cerevisiae;* specific mutations in these factors give rise to suppressor of upstream initiation codon (Sui-) phenotypes and their suppressors (Ssu-) (61). To ask if between-species variability in start codon preference is linked to these initiation factors, we generated multiple sequence alignments of their homologs in fungi.

Translation initiation factor eIF1 shows striking sequence variation across fungi, notably at multiple *Cryptococcus-s*pecific sequence insertions that result in a 159-aa protein substantially larger than the 108-aa S. *cerevisiae* homolog (Figure 9A). Variation in eIF1 occurs at and around positions known to modulate start codon selection inS. *cerevisiae* (61). For instance, a T15A substitution increases fidelity in SceIF1 (61), and an analogous T15A substitution is present in eIF1s from *Neurospora* and other filamentous fungi, while both *Cryptococcus* homologs have the T15V substitution. The three fungi that tend not to use alternative AUG start codons in the regulation of proteome diversity, S. *cerevisiae, C. albicans, andS. pombe*, all have a threonine residue at position 15. Variation in fungal eIF1 extends far beyond this N-terminal region: similar patterns of sequence diversity occur at the positions E48, L51, D61 that have been shown to increase fidelity in SceIF1 (61). By contrast, positions K56, K59, D83, Q84, at which mutations have been shown to reduce fidelity in SceIF1 (61), are highly conserved in fungi.

**Figure 9.**
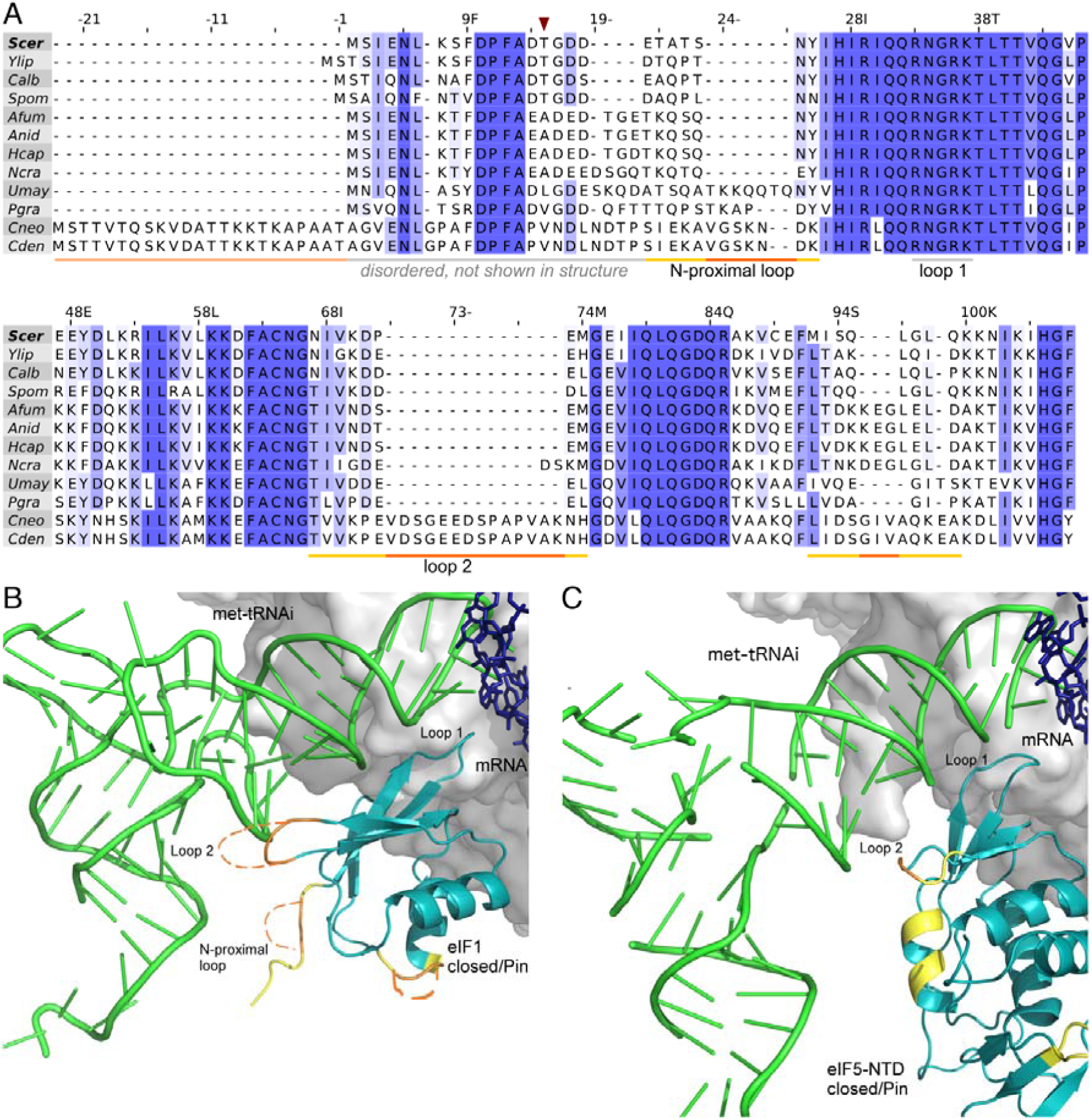
Eukaryotic translation initiation factor 1 is highly variable across fungi. A, Multiple sequence alignment of translation initiation factor eIF1 from 12 fungi, numbered asS. *cerevisiae (Seer*, top line). *Cryptococcus* insertions are indicated in orange, and surrounding variable residues in yellow. The N-terminal extension in *Cryptococcus* elF1, that is predicted disorded, is shown in pale orange, and T15 residue with dark red arrow. B, Structural predictions of insertions (orange) and non-conserved neighborhoods (yellow) in *Cryptococcus* eIF1 mapped onto the closed pre-initiation complex of S. *cerevisiae/K.lactis* (PDB:3J81, Hussein 2015). eIF1 (teal) and Met-tRNAi (green) in closed conformation, shown with synthetic mRNA sequence (pink), and eIF2 (pale pink) and ribosomal subunit surface (greys) in background. Approximate ribosomal contacts are shown as grey background surface and eIF2-alpha subunit is shown as pale pink sticks. B, Structural predictions of variations in *Cryptococcus* eIF5 mapped on toS. *cerevisiae* PIC (PDB:6FYX, (27)). Multiple sequence alignment of eIF5 is shown in figure S9.1A.

We next tested how the translation pre-initiation complex could be affected by the insertions in *Cryptococcus* eIF1 using published structures of the S. *cerevisiae/K. lactis* “Pin” complex engaged in the act of AUG selection (62). We found that the insertions in eIF1 are facing either the methionine initiator tRNA (RNAi) or the solvent-exposed side (Figure 9B). The N-terminal insertion is not visible in the structure, but could be close to the acceptor arm of tRNAi. The N-proximal loop insertion of CneIF1 extends from the SceIF1 sequence (18-DETATSNY-25) that contacts the acceptor arm of tRNAi. The CneIF1 insertion in loop 2 extends the SceIF1 loop 2 (70-KDPEMGE-76) that contacts the D-loop of tRNAi; substitutions D71A/R and M74A/R increase the charge of SceIF1 loop 2 and increase initiation at UUG codons and weak AUG codons (63). CneIF1 loop 2 has substitutions at both these functionally important sites, and is extended by a further 14 hydrophobic and negative residues. The last insertion in CneIF1 extends a loop facing the solvent-exposed surface of SceIF1. Collectively, this shows that there are likely major differences in the eIF1-tRNAi interaction surface in *Cryptococcus* relative to other fungi, an interaction critical for start codon selection (63).

The N-terminal domain of eIF5 (eIF5-NTD) replaces eIF1 upon start codon recognition, and we found between-species variation in CneIF5 at tRNAi interaction surfaces corresponding to variability in CneIF1 (Figure 9C, S9.1A). SceIF5 Lys71 and Arg73 in loop 2 make more favourable contacts with the tRNAi than the corresponding residues of SceIF1, so that the shorter loop 2 of SceIF5 may allow the tRNAi to tilt more towards the 40S subunit (27). Although Arg73 is conserved across fungi, Lys71 is absent in CneIF5 loop 2 (67-SMAN-70), which is two amino acids shorter than SceIF5 loop 2 (66-SISVDK-71). Collectively, the longer loop 2 of CneIF1 and the shorter loop 2 of CneIF5 suggest that the conformational changes accompanying start codon recognition may be more exaggerated in *Cryptococcus*, providing a mechanistic hypothesis for stronger genomic patterns of start codon recognition. Fungal eIF1A homologs also diverge from SceIF1A at regions that modulate translation initiation fidelity (Figure S9.1B), for example the N-terminal element DSDGP (61). The *Cryptococcus* eIF1A C-terminus is diverged from all other fungi at SceIF1A positions 110-120, and along with other basidiomycetes lacks a loop at SceIF1A positions 135-149. This C-terminal region of SceIF1A contributes to pre-initiation complex assembly and binds eIF5B (64) and eIF5 (65), and domain deletions or local alanine substitutions reduce fidelity of translation start site selection (61, 64, 66).

Thus, although structural analysis of the Cryptococcal initiation complex will be required for a detailed mechanistic understanding, our initial analysis suggests that sequence variability in fungal eIFs could plausibly account for differences in start codon selection between different species.

## DISCUSSION

Our annotation of transcript structure and translation in two pathogenic *Cryptococcus* species and our analysis of published data from other species show that start codon context has a major effect on protein production, regulation, diversity, and localization in the fungal kingdom. As such this work represents a useful resource for the field. While the genomewide effect of start codon context is weak in S. *cerevisiae* (20), we find that other fungi, from *Neurospora* to *Cryptococcus*, use start codon context to regulate translation initiation to a far greater extent. These fungi have long and AUG-rich TLs, and more information-rich and functionally important Kozak sequences. Further, *Cryptococcus* and *Neurospora* display extensive evidence of leaky scanning of weak AUG codons that is used for regulation by upstream ORFs and to generate alternate N-terminal isoforms with different subcellular localization.

### Widespread leaky scanning controlled by start codon context in *C. neoformans*

Translation initiation regulation can be enabled by start codons that are imperfectly used, so that scanning pre-initiation complexes can leak past them. According to the scanning model of translation initiation, a “perfect” strong start codon would prevent this by capturing all the scanning PICs, and leave none for regulatable downstream initiation. For example, the downstream out-of-frame ORF of *Cryptococcus* eIF1 is likely to be translated only by PICs that leak past the annotated AUG. The alternative second in-frame AUG of dual-localized proteins is also initiated only by PICs that have leaked past the initial AUG. Our data show this leakiness-driven dual-localization is common in *Cryptococcus*, in addition to being conserved across eukaryotes in gene classes such as tRNA synthetases. Our data also argue that AUGs that are proximal to the 5’ cap, or that have poor sequence context, are commonly leaked past in *Cryptococcus*, as shown previously in studies of yeast (67) and mammals (17, 68). We note that leakiness-driven translation regulation is not the only mechanism regulating alternative translation from a single mRNA and is distinct from those that depend on either blocking scanning, or on recycling of post-termination ribosomes such as in the case of S. *cerevisiae GCN4* (37).

### Functional role of start codon context varies across the fungal kingdom

*Cryptococcus* and *Neurospora* have long TLs that are AUG-rich, and extended start codon context sequences that suggest a higher ability to discriminate against poor-context AUGs. Several lines of evidence argue that efficiency with which upstream AUGs capture initiation complexes is determined by the AUG sequence context. The most spectacular examples of uORF-associated translation repression in *Cryptococcus* are associated with good-context uAUGs with high ribosome occupancy. However, such strong-context high-occupancy uAUGs are rare. In *Cryptococcus* and *Neurospora*, the leakiness of potential AUG translation start sites is also extensively used to diversity the proteome by alternative N-terminal formation. In comparison, S. *cerevisiae, S. pombe* and *C. albicans* appear to be less efficient in discriminating AUGs based on their sequence context. S. *cerevisiae* has minimized the possibility of regulation of gene expression by uORFs: it has unusually short TLs, these TLs are unusually AUG-poor, uAUGs tend to have poor context, and there is no statistical association between uAUG score and translation efficiency of the main ORF. Reporter gene studies (21, 22) and classic examples such as *GCN4* show that uAUGs can repress translation in S. *cerevisiae*, but genome-wide analysis show that this is rare during exponential growth in rich media (Fig S5.1). Recent work on meiosis (69) and stress (70) shows that 5’-extended transcript leaders that contain repressive uAUGs (“long undecoded transcript isoforms”) are more common during alternative growth conditions for this yeast. Moreover, in S. *cerevisiae*, near-cognate codons appear to be more common starts for alternative N-terminal formation (71). This suggests that leaky scanning from near-cognate codons, more than from AUGs, might be an important mode of regulation in S. *cerevisiae*. The situation is different in S. *pombe*, which has long AUG-rich TLs but is depleted for downstream in-frame AUGs. Consequently, uAUGs globally repress gene expression, but do not appear to regulate alternative protein production through alternative AUG start codons. We speculate that the comparatively uninformative Kozak context in S. *pombe* might be variable enough to regulate translation initiation rate but not proteome diversity.

We found that multiple near-cognate start codons are used for leaky initiation in *Cryptococcus:* ACG for the mitochondrial isoform of LeuRS, AUU for the mitochondrial isoform of ArgRS, and the upstream CUG in eIF5. Further work will be needed to quantify the extent of near-cognate start codon usage in *Cryptococcus* in different growth conditions and compare it to other organisms (14, 72).

### Leaky scanning through weak AUGs could regulate the mitochondrial proteome

We computationally predicted dozens of dual-localized proteins with alternative start codons that confer an N-terminal mitochondrial targeting sequence in their longest isoform. We did not identify enrichment of proteins with predicted dual-localization in the cytoplasm and in the nucleus, or with a signal peptide followed by an alternative start codon (data not shown). Thus, increasing the efficiency of weak-context to strong-context translation initiation would predominantly upregulate a regulon consisting of the mitochondrial isoforms of dozens of proteins.

Mechanisms to control initiation efficiency of a mitochondrial-localized regulon could include intracellular magnesium concentration (73), variations in availability or modification status of shared initiation factors, variations of the ratio of mitochondrial volume to intracellular volume (74), or specialized factors to promote initiation specifically of mitochondrial isoforms with their specialized start codon context. Nakagawa et al (75) previously suggested that distinct Kozak contexts might be recognized by different molecular mechanisms.

One candidate mechanism involves the translation initiation factor 3 complex, which has a role in regulating the translation initiation of mitochondrial-localized proteins across eukaryotes. In 5. *pombe*, subunits eIF3d/e promote the synthesis of mitochondrial electron transfer chain proteins through a TL-mediated mechanism (76). InS. *cerevisiae* and *Dictyostelium discoideum* the conserved eIF3-associated Clu1/CluA protein affects mitochondrial morphology (77), and the mammalian homolog CLUH binds and regulates mRNAs of nuclear-encoded mitochondrial proteins (78, 79). Metazoans have 12 stably-associated subunits of eIF3, which are conserved in most fungi including *N. crassa* (80), *Cryptococcus*, and the Saccharomycetale yeast *Yarrowia lipolytica* (Table S7). Interestingly, species that tend not to use alternate AUG codons for dual-localization have lost eIF3 subunits: eIF3d/e/k/l/m are lost in *C. albicans*, and additionally eIF3f/h in the related S. *cerevisiae; S. pombe* has independently lost eIF3k/I (Table S7; (51)). Further work will be needed to investigate the role of eIF3 in regulating mitochondrial- and dual-localized proteins in the fungal kingdom.

### Evolutionary plasticity of translational initiation in the fungal kingdom

Selection on genome compaction in unicellular yeasts, which has independently led to gene loss and high gene density in multiple lineages of yeast, could lead to shorter TLs. However, *Saccharomyces, Schizosaccharomyces*, and *Cryptococcus* have all independently evolved yeast lifestyles with compact genomes, yet their average TL lengths differ three-fold. Mutations in gene expression machinery, such as the variation in eIF1 noted above, would alter selective pressure on start codon context, and thus uAUG density. Cells have multiple redundant quality control mechanisms, and flexible protein production through leaky scanning could be buffered by such mechanisms enabling their evolution. Key control mechanisms acting on mRNA, such as RNAi and polyuridylation, have been lost in fungal lineages such as *Saccharomyces*, which might explain their more ‘hard-wired’ mechanism of translation initiation.

Unexpectedly, highly conserved core translation initiation factors, such as eIF1, have distinctive sequence inserts in *Cryptococcus* that are not shared even by basidiomycetes such as *Puccinia* and *Ustilago*. One possibility is genetic conflict, as genetic parasites hijack the gene expression machinery (81). Thus, the unique aspects of the *Cryptococcus* translation initiation machinery could have arisen from a past genetic conflict in which rapid evolution of initiation factors in an ancestor enabled evasion of a genomic parasite (e.g. a mycovirus) that would otherwise hijack initiation.

## Supporting information

Table S1: Sequencing and annotation numbers.

Table S2: Differential expression results in Cryptococcus deneoformans upf1??.

Table S3: Cytoplasmic ribosomal proteins in 6 fungal species.

Table S4: Genes with score d1AUG ??? aAUG > 0.1, n = 167 (Excel table).

Table S5: List of aaRS in Cryptococcus and select fungi (Excel table).

Table S6: Initiation contexts of annotated and downstream AUGs in 9 Cryptococcus aaRSs

Table S7: Initiation factor 3 components in 12 fungal species. (Excel table)

Table S8: rRNA subtraction oligos used in ribosome profiling.

## Data Availability

Raw and summarized sequencing data are available on GEO under accession numbers <u>GSE133695</u> (RNA-seq, TSS-seq, PAS-seq) and GSE133125 (ribosome profiling and matched RNA-seq).

## Acknowledgments

We thank members of the Wallace, Janbon, and Madhani labs for helpful discussions and comments on the manuscript. We thank Juan Mata for sharing intermediate data related to (Duncan and Mata 2017). We are grateful to J. Weissman (UCSF) for advice on ribosome profiling. Work in the Madhani lab is supported by grants from the US National Institutes of Health. H.D.M. is an Investigator of the Chan-Zuckerberg Biohub. E.W.J.W. is a Sir Henry Dale Fellow, supported by a Sir Henry Dale Fellowship jointly funded by the Wellcome Trust and the Royal Society (Grant Number 208779/Z/17/Z). L.T. is supported by a Wellcome-University of Edinburgh ISSF3 award.

## Material and Methods

### DNA and RNA purification, sequencing library preparation

*C. neoformans* strain H99 and *C. deneoformans* strain JEC21 and were grown in YPD at 30°C or 37°C under agitation up to exponential or early stationary phase as previously described (32). Briefly, early stationary phase was obtained after 18 h of growth (final OD_600_= 15) starting from at OD_600_ =0.5. *C. deneoformans* strain NE579 (*upf1Δ*) (33) was grown in YPD at 30°C under agitation in exponential phase. Each *Cryptococcus* cell preparation was spiked in with one tenth (OD/OD) of S. *cerevisiae* strain FY834 (82) cells grown in YPD at 30°C in stationary phase. Cells were washed, snap frozen and used to prepare RNA and total DNA samples as previously described (32). Each condition was used to prepare biological triplicate samples.

For RNA-Seq, strand-specific, paired-end cDNA libraries were prepared from 10 μg of total RNA by polyA selection using the TruSeq Stranded mRNA kit (Illumina) according to manufacturer’s instructions. cDNA fragments of ~400 bp were purified from each library and confirmed for quality by Bioanalyzer (Agilent). DNA-Seq libraries were prepared using the kit TruSeq DNA PCR-Free (Illumina). Then, 100 bases were sequenced from both ends using an Illumina HiSeq2500 instrument according to the manufacturer’s instructions (Illumina).

TSS-Seq libraries preparations were performed starting with 75 μg of total RNA as previously described (34) replacing the TAP enzyme by the Cap-clip Pyrophosphatase Acid (TebuBio). For each *Cryptococcus* species we also constructed a control “no decap” library. Briefly, for these control libraries, poly A RNAs were purified from 75 μg of RNA from *Cryptococcus* and 75 μg of RNAS. *cerevisiae* before being dephosphorylated using Antarctic phosphatase. Then, S. *cerevisiae* RNAs and one half of the RNAs extracted from *Cryptococcus* were treated with Cap-clip Pyrophosphatase Acid enzyme. The second half of *Cryptococcus* RNAs was mock treated. Each half of Cap-clip Pyrophosphatase Acid *Cryptococcus* RNA samples was mixed with the same quantity of S. *cerevisiae* Cop-clip Pyrophosphatase Acid treated RNAs. The subsequent steps of the library preparation were identical to the published protocol (34). Single 50 bases single end reads were obtained using an an Illumina HiSeq2500 instrument according to the manufacturer’s instructions (Illumina).

For QuantSeq 3’mRNA-Seq preparation we followed the manufacturer instructions (Lexogen GmbH, Austria). 100 base single end reads were obtained using an Illumina HiSeq2000 instrument according to the manufacturer’s instructions (Illumina).

### Sequencing data analyses

For TSS analysis we kept only the reads containing both the oligo 3665 (AGATCGGAAGAGCACACGTCTGAAC) and the 11NCGCCGCGNNN tag (34). These sequences were removed and the trimmed reads were mapped to the Cryptococcus genome and S. cerevisiae genomes using Bowtie2 and Tophat2 (83). Their 5’ extremities were considered as potential TSSs. For each condition we kept only the positions that were present in all three replicates. Their coverage was normalized using the normalization factor used for spiked in RNA-Seq. TSS positions were then clustered per condition. As most of the observed TSS sites appeared as clusters, we grouped them into clusters by allowing an optimal maximum intracluster distance (at 50 nt) between sites as previously used (34). We then removed the false TSS clusters using the “no-cap” data keeping the clusters i for which

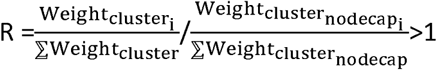

Similarly, QuantSeq 3’mRNA-Seq reads containing both the Sequencing and indexing primers (Lexogen) were sorted. The reads were then cleaned using cutadapt/1.18 (84) and trimmed for polyA sequence in their 3’end. PolyA untrimmed and trimmed reads were mapped to the adapted *Cryptococcus* and to the S. *cerevisiae* genomes with Tophat2 (83) with the same setting as for RNA-Seq. To eliminate the polyadenylated reads corresponding to genomic polyA stretches, we considered only the reads that aligned to the genomes after polyA trimming but not before the trimming. The 3’end position of these reads were considered as potential PAS. As for the TSS, for each condition we kept only the positions that were present in all three replicates. Similarly, the PAS dataset was normalized using the spike in normalization factor and the PAS positions were clustered using the same strategies.

### Ribosome profiling and matched mRNA-seq

Ribosome profiling was performed on both *C. neoformans* H99 and *C. deneoformans* JEC21, two biological replicates of WT-H99 and one replicate each of H99 *ago1Δ* and H99 *gwo1Δ* strains from (30), and one replicate each of WT-JEC21 and JEC21 *ago1Δ* There was negligible differential expression detected between these deletions and their background strains, so in our analyses we treat the deletion strains as biological replicates.

Cells were grown to exponential phase in 750 mL of YPAD with shaking at 30°C. 100 ug/ml cycloheximide (Sigma) (dissolved in 100% ethanol) was added to the culture and incubated for 2 minutes. 50 mL of the culture was withdrawn for performing RNA-Seq in parallel. Cells were then pelleted, resuspended in 5ml of lysis buffer (50mM Tris-HCl pH. 7.5, 150mM NaCl, lOmM MgCl2, 5mM DTT, 0.5% Triton and 100ug/mL cyclohexamide) and snap frozen. Lysis, clarification, RNasel digestion, sucrose gradient separation and monosome isolation was performed as previously described (36).

Ribosome protected fragments were isolated from the monosome fraction using hot phenol. 150ug of the total RNA extracted from the 50 ml of culture in parallel was polyA selected using the Dynabeads mRNA purification kit (Thermo Fisher Scientific) and digested using freshly made fragmentation buffer (100mM NaCO3 pH. 9.2 and 2mM EDTA) for exactly 20 mins.

RNA was resolved on a 15% TBU gel. A gel slab corresponding to 28-34 nt was excised for footprint samples and around 50 nt for mRNA samples, then eluted and precipitated. Sequencing libraries were generated from the RNA fragments as described in Dunn et al. with the following modifications (85). cDNA was synthesized using primer oCJll (Table S8). Two rounds of subtractive hybridization for rRNA removal was done using oligos rasl-8 listed in Table S8. After circularization Illumina adaptors were added through 9 cycles of PCR. Libraries were sequenced on a HiSeq 2500 (Illumina).

### Ribosome profiling data analysis

Ribosome profiling and matched RNA-seq reads were demultiplexed on BaseSpace (Illumina) and then analyzed essentially with the RiboViz pipeline v.1.1.0 (86). In brief, sequencing adapters were removed with cutadapt (84), and then reads aligned to rRNA were removed by alignment with hisat2 (87). Cleaned non-rRNA reads were aligned to (spliced) transcripts with hisat2 (87), sorted and indexed with samtools (88), and then quantified on annotated ORFs with bedtools (89), followed by calculation of transcripts per million (TPM) and quality control with R (90) scripts included in RiboViz. The cleaned non-rRNA reads were also aligned to the genome with hisat2, and processed analogously, then used to generate figures of genome alignments using ggplot2 (91) in R (90).

### Data analysis and visualization

Data analysis and visualization were scripted in R (90), making extensive use of dplyr (92), ggplot2 (91), and cowplot (93). Sequence logos were prepared in ggseqlogo (94). Analysis of differential expression for *upf1Δ* data was performed in DeSeq2 (95). Figures were assembled and annotated in Inkscape v0.92 (https://inkscape.org).

Protein sequences were aligned using muscle (96), with default parameters for protein sequences and 100 iterations. Phylogenetic trees were constructed using ClustalW2 tool v2.1 (97) by using the neighbor-joining method with 1000 bootstrap trial replications. Structural figures were prepared in PyMOL (Schrodinger).

### External datasets

*N. crassa* (strain OR74A) ribosome profiling data from ((14), GEO:GSE97717) was used to generate highly-translated genes, and ribosome profiling and RNA-seq data from ((98), GEO: GSE71032) used to estimated TE. In both cases, we estimated TPMs using the RiboViz pipeline as above, using the NC12 genome annotation downloaded from EnsembIGenomes (99). TL sequences were also obtained from NC12.

*S. pombe* (strain 972h) ribosome profiling and RNA-seq data are from (100), and the authors provided us with a table of RPKMs for all replicates as described. Genome sequence and annotation ASM294v2, including TL annotation, were downloaded from EnsembIGenomes (99).

*C. albicans* (strain SC5314) ribosome profiling and RNA-seq data are from (101), GEO:GSE52236), processed with the RiboViz pipeline as above using the assembly 22 of the strain SC5414 genome annotation from CGD (102).

*S. cerevisiae* (strain S288C/BY4741) highly-translated genes use the RPKM table from ((103), GEO:GSE59573), and highly-expressed genes use (104). For TE estimates, we used matched ribosome profiling and RNA-seq estimates from (105), although we did not use this for the list of highly translated genes because near-duplicate paralogous ribosomal protein genes were not present in the dataset, which thus omits a substantial fraction of highly-translated genes. TL sequences were downloaded from SGD (106)

Protein homolog lists were assembled with OrthoDB (47) and PANTHERdb (48), with reference to FungiDB (107). The list of cytoplasmic ribosomal proteins was assembled in 5. *cerevisiae* based on (108) with help from SGD (106), extended to other fungi with PANTHERdb (47), and manually curated.

---

**Figure S2.1, related to figure 2:**
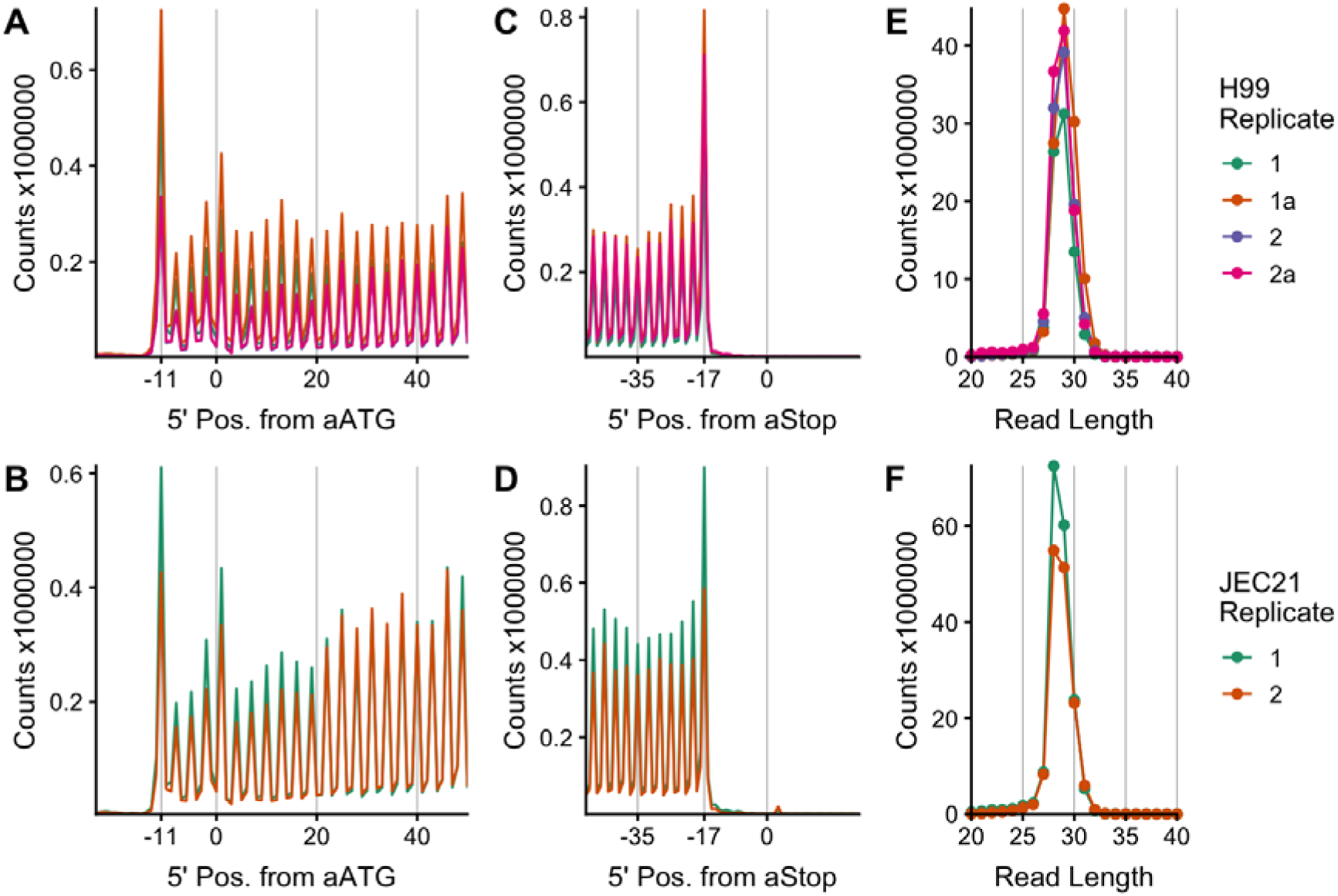
Ribosome profiling data passes quality control metrics. Metagene profiles of mapped 5’ ends of ribosome-protected fragment counts at the 5’ end (A,B) and 3’ end (C,D) of ORFs, showing 3-nucleotide periodicity indicative of active translation starting at the annotated start codon and ending at the annotated stop codon. Ribosome protected fragment length is of a consistent length with other studies (E,F). Top row is data from 4 replicates of *C. neoformans* H99, bottom row form 2 replicates of *C. deneoformans* JEC21. These figures were made using RiboViz (86).

**Figure S2.2, related to figure 2:**
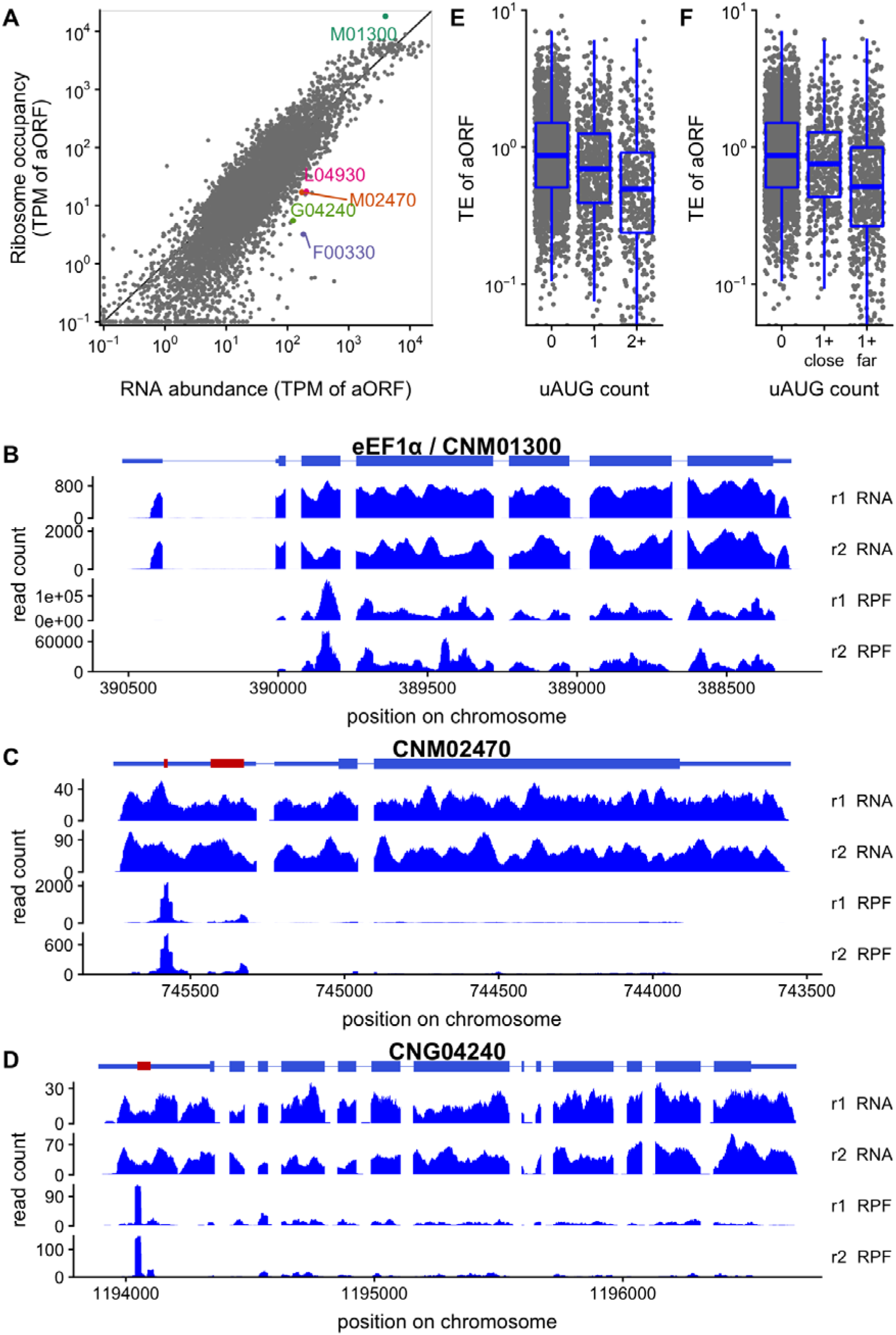
Upstream AUGs repress translation in *C. deneoformans*. A, translation regulation of annotated ORFs (aORFs) in *C. deneoformans* JEC21 growing exponentially in YPD at 30°C. Ribosome occupancy is plotted against the RNA abundance, both calculated in transcripts per million (TPM) on the aORF. B, uAUGs are associated with lower translation efficiency (TE) of annotated ORFs, measured as the ratio of ribosome occupancy to RNA-seq reads. C, only uAUGs far from the transcription start site are associated with low TE. A gene is in the “1+ far” category if it has at least one uAUG more than 20nt from the TSS, “1+ close” if all uAUGs are within 20nt of the TSS. D-F, Examples of ribosome occupancy profiles along select RNAs highlighted in A (others are shown in Fig S2B). D, Translation elongation factor eEF2/CNM01300 (CNAG_06125 homolog) has high ribosome occupancy in the annotated ORF. Translationally repressed mRNAs CNM02470 (CNAG_06246 homolog, E) and CNG04240 (CNAG_03140 homolog, F) have high ribosome occupancy in uORFs in the transcript leader (red), and low ribosome occupancy in the aORF. Only the first of 5 uORFs in CNG04240 is shown, and only transcript isoform tOl is shown, excluding the annotated TL intron in isoform t02.

**Figure S2.3, related to figure 2:**
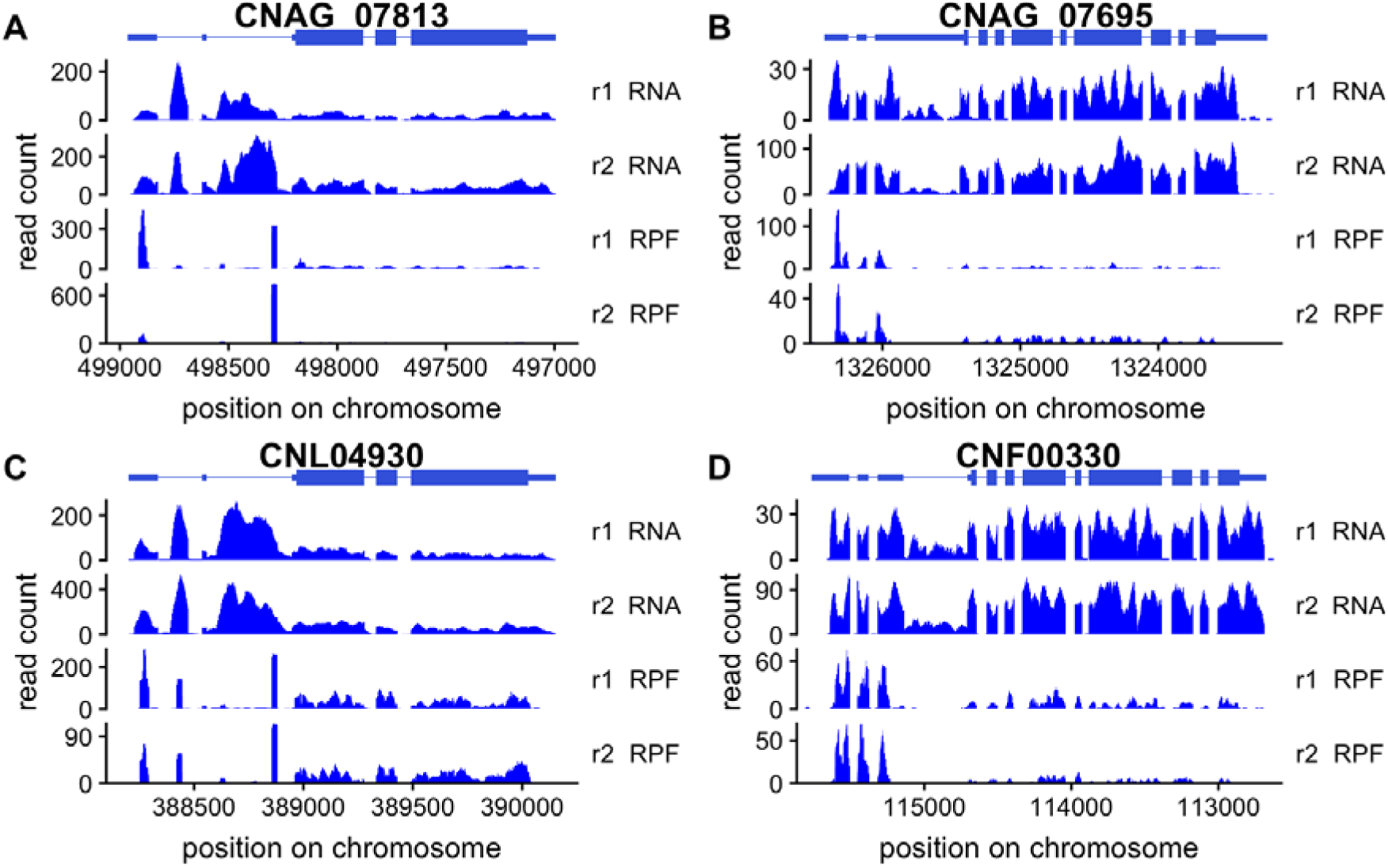
Further examples of upstream AUG and 5’-end regulation in *C. neoformans* and *C. deneoformans*. CNAG_07813 (A) and CNL04930 (C) are paralogs, and in addition to an upstream ORF with ribosome occupancy, they have an intronically encoded non-coding RNA in the TL. CNAG_07695 (B) and CNF00330 (D) are paralogs, and in addition to an upstream ORF with ribosome occupancy, they have an alternatively-spliced intron in the TL that is not occupied by ribosomes.

**Figure S3.1, related to Figure 3:**
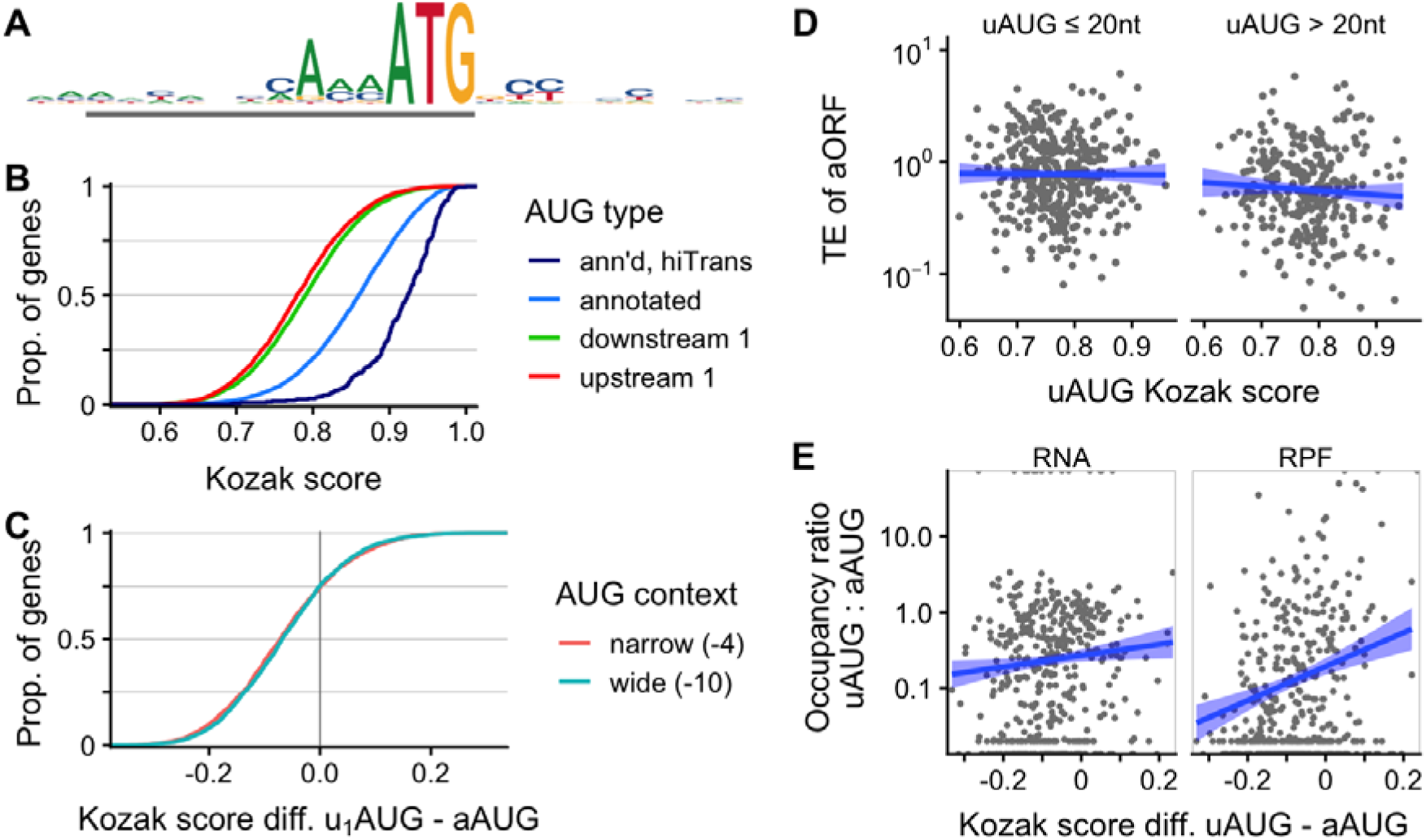
AUG sequence context is associated with translation in *C. deneoformans*. As for Fig 3, but with data from *C. deneoformans* JEC21.

**Figure S4.1, related to Figure 4:**
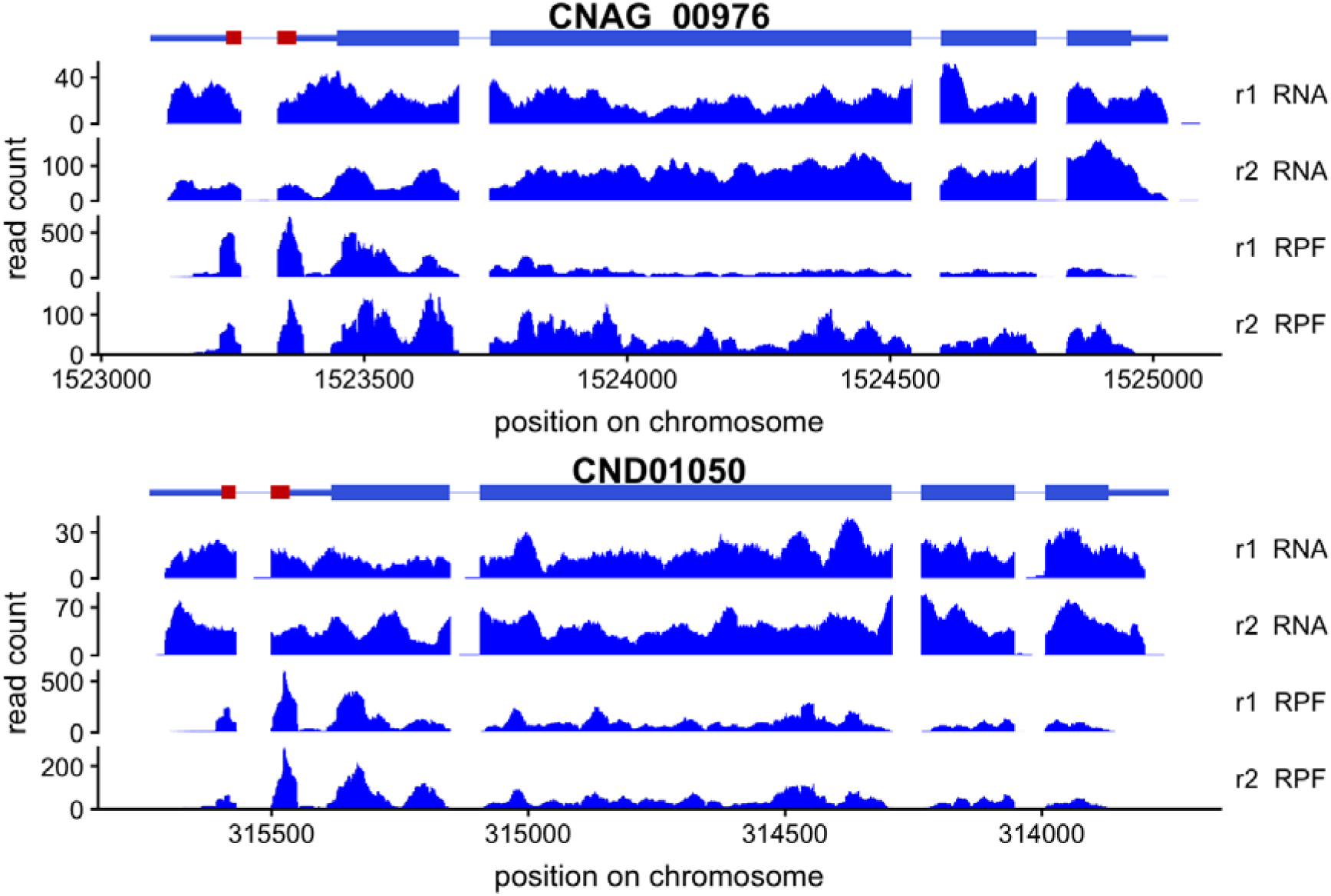
Carbamoyl-phosphate synthase CPA1 homologs have a conserved uORF that is occupied by ribosomes in *C. neoformans* (A) and *C. deneoformans* (B).

**Figure S5.1, related to figure 5:**
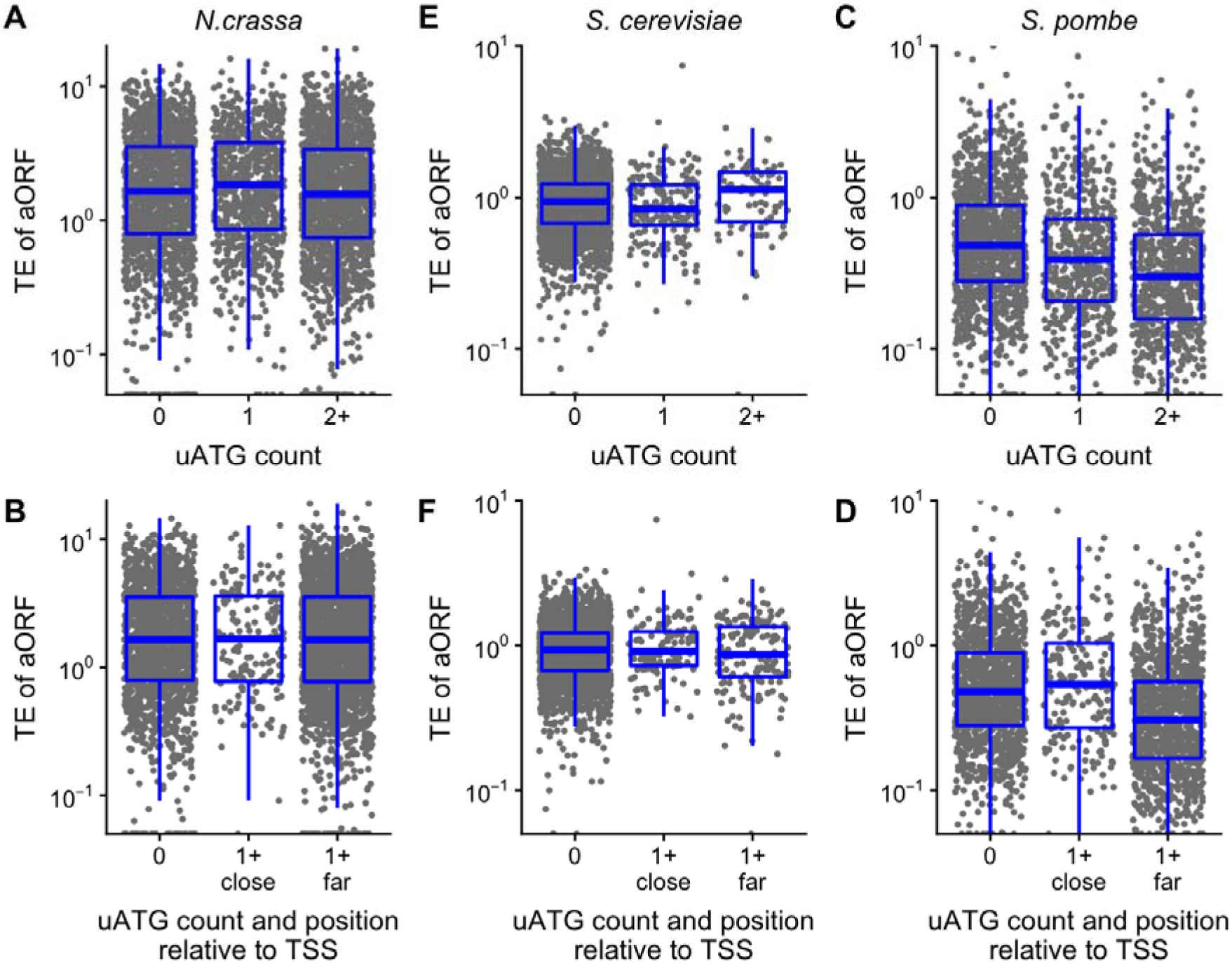
Effect of uATGs on translational efficiency in *N. crassa, S. pombe*, and S. *cerevisiae*.

**Figure S6.1, related to Figure 6:**
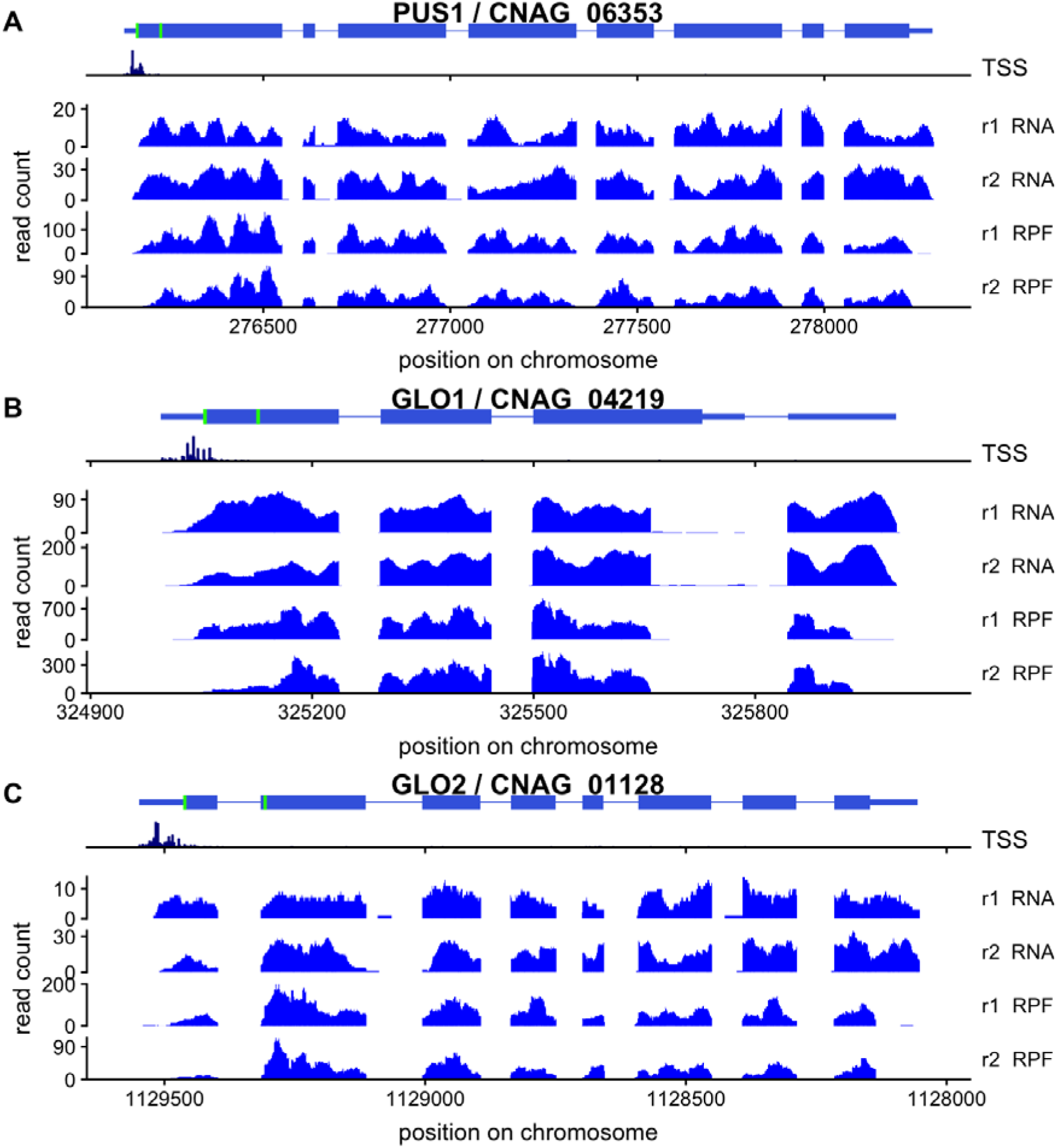
Ribosome profiles of *C. neoformans* genes with predicted dual-localization specified by alternative N-termini.

**Figure S7.1, related to Figure 7:**
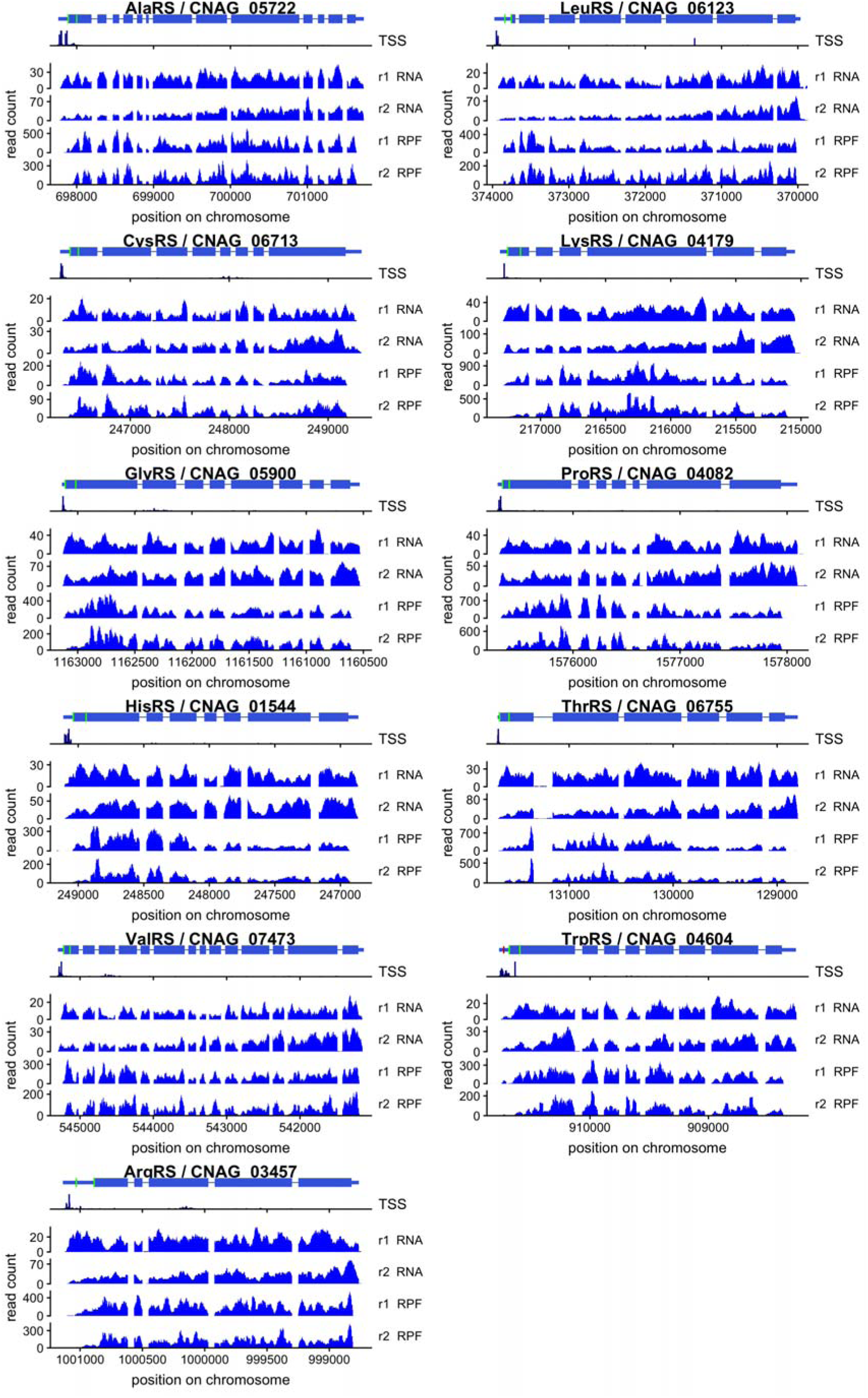
Ribosome profiles along the 11*C. neoformans* aaRS genes with predicted dual-localization. Predicted start codons are shown in green, and the uORF of TrpRS in dark red.

**Figure S8, related to Figure 8:**
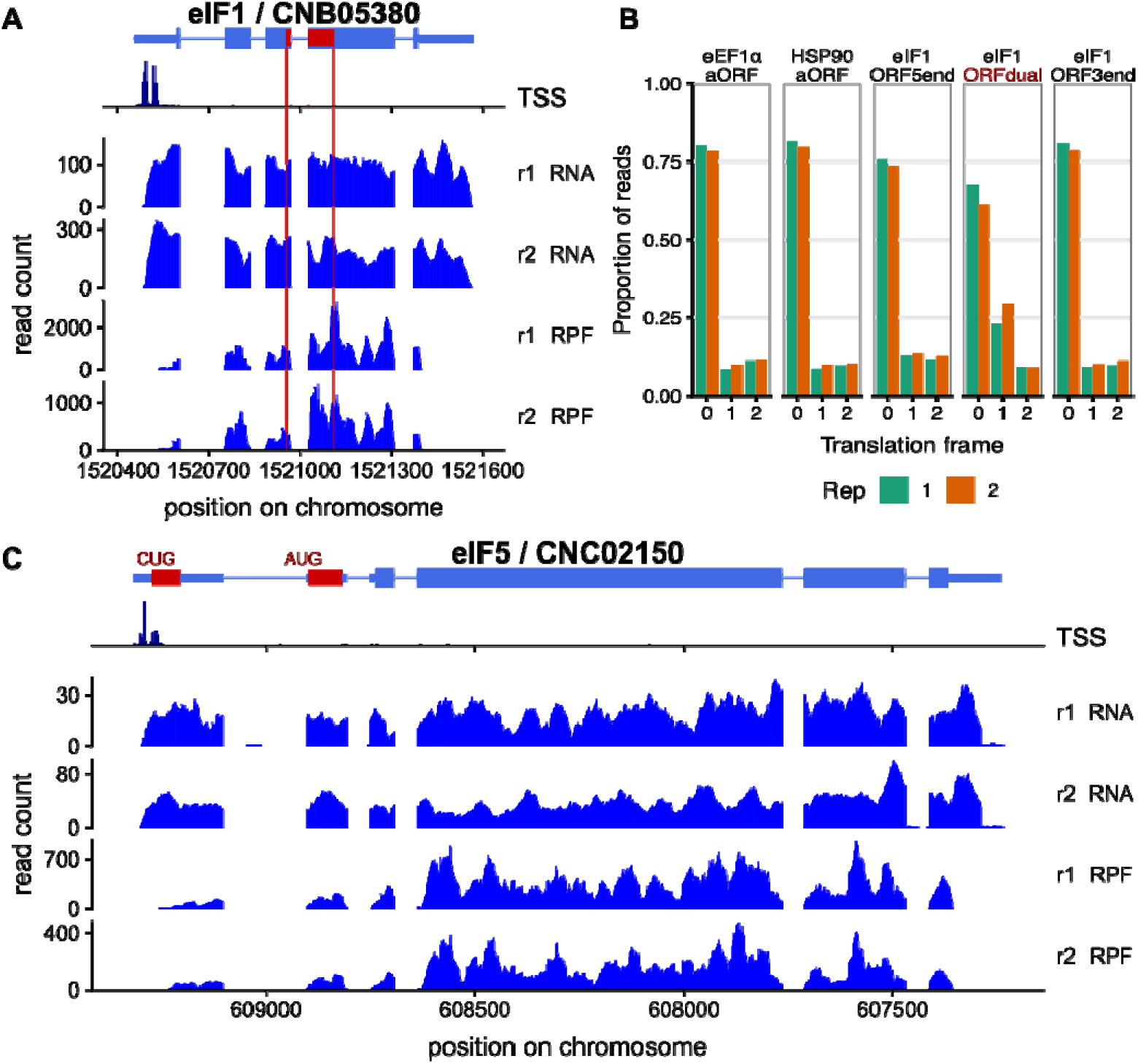
Translation initiation factors eIF1 and eIF5 are regulated by alternate start codon usage in *C. deneoformans*. See figure 8 for legend.

**Figure S9.1, related to Figure 9:**
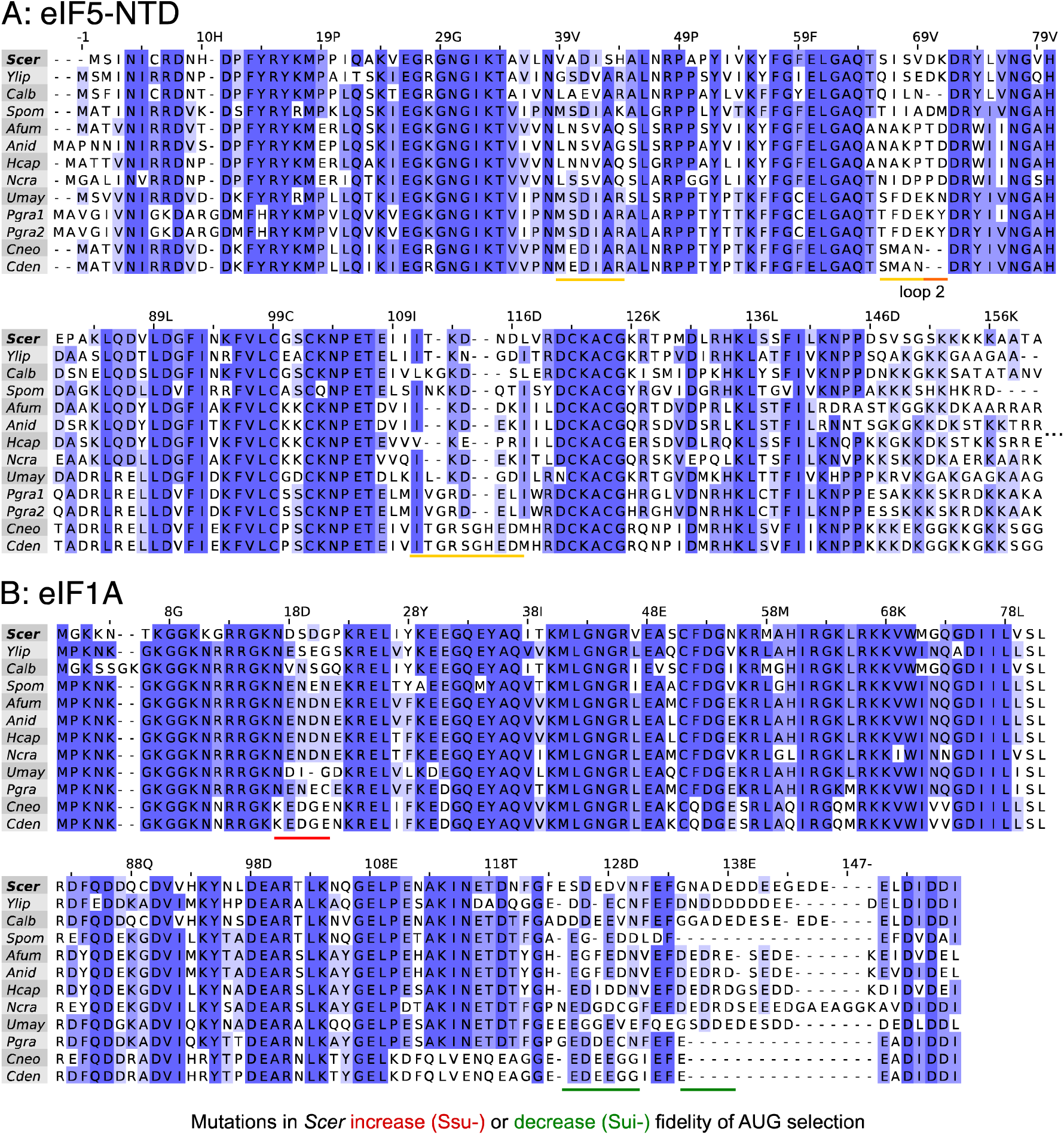
multiple sequence alignments of eIF5-NTD and eIF1A from select fungi. Sequences are numbered according to the S. *cerevisiae* homologs.

**Table S1: Sequencing and annotation numbers (Excel table).**

**Table S2: Differential expression results in Cryptococcus deneoformans upf1Δ.**

**Table S3: Cytoplasmic ribosomal proteins in 6 fungal species.**

**Table S4: Genes with score d_1_AUG - aAUG > 0.1, n = 167 (Excel table).**

**Table S5: List of aaRS in Cryptococcus and select fungi (Excel table).**

**Table S6: Initiation contexts of annotated and downstream AUGs in 9 Cryptococcus aaRSs**

**Table S7: Initiation factor 3 components in 12 fungal species. (Excel table)** We show homologs of all human eIF3 components eIF3a-eIF3m, assembled mostly from PANTHERdb (51). Note that the *C. neoformans* homolog *PRT1* is expressed in two nonidentical paralogs at the mating-type locus, so does not have a systematic ORF name.

**Table S8: rRNA subtraction oligos used in ribosome profiling.**

